# Integrating Biometric, Metabolic, and Photochemical Traits to Identify Candidate Physiological Indicators of Salinity Tolerance and Forage Quality in Alfalfa Cultivars

**DOI:** 10.64898/2026.11.18.767659

**Authors:** Giuliano Barbieri, Mónica Cornacchione, Dario Fernando Luna, Paolo Annicchiarico, Marianela S. Rodriguez

**Affiliations:** Unidad de Estudios Agropecuarios (UDEA- INTA-CONICET) Camino 60 cuadras km 5.5 X5020ICA, Córdoba, Argentina; Instituto Nacional de Tecnología Agropecuaria (INTA), Estación Experimental Santiago del Estero, Jujuy 850, Santiago del Estero, 4200, Argentina; Instituto de Fisiología y Recursos Genéticos Vegetales, Ing Agr Victorio Segundo Trippi (IFRGV), Centro de Investigaciones Agropecuarias (CIAP), Instituto Nacional de Tecnología Agropecuaria (INTA), Camino 60 cuadras km 5.5, X5020ICA, Córdoba, Argentina; Consiglio per la Ricerca in Agricoltura e l’Analisi dell’Economia Agraria (CREA), Centro di Ricerca Zootecnia e Acquacoltura, Lodi, Italy

**Keywords:** alfalfa, chlorophyll fluorescence, ion homeostasis, photochemical acclimation, salinity tolerance, forage quality

## Abstract

Salinity limits alfalfa (*Medicago sativa* L.) productivity, yet integrative physiological markers of cultivar tolerance remain poorly defined. This study aimed to identify candidate physiological indicators of salt tolerance by integrating growth, metabolic, and time-resolved photosynthetic traits across diverse alfalfa cultivars.
We exposed nine alfalfa cultivars to a 200 mM NaCl hydroponic shock and monitored their physiological responses over 14 days. We assessed shoot and root growth, leaf Na^+^/K^+^ homeostasis, carbohydrate partitioning, dynamic chlorophyll a fluorescence (ChlF), and NIRS-based forage quality.
Salinity elicited cultivar-dependent responses. Tolerant cultivars-maintained shoot biomass, restricted foliar Na^+^ accumulation, preserved leaf starch reserves and progressively recovered Phi2 and linear electron flow (LEF) following initial declines. In contrast, less tolerant cultivars showed persistent growth inhibition, reduced leaf starch, and persistent photochemical impairment. At 7 and 14 dpt, PI_total_ and Fv’/Fm’ correlated positively with final biomass, whereas PhiNPQ and PhiNO showed negative correlations. Notably, the association between regulated energy dissipation (PhiNPQ) and biomass retention inverted over time—shifting from an early photoprotective mechanism at 1 dpt to a sign of chronic photoinhibitory strain by 7 and 14 dpt. These responses were accompanied by cultivar-specific changes in Na^+^/K^+^ balance, soluble sugars, starch, and forage quality.
Tracking time-resolved PSII photochemistry reveals that maintaining *PI*_total_ and Fv’/Fm’ under saline stress offers an accurate, non-invasive proxy for evaluating cultivar resilience. Their integration into high-throughput phenotyping and breeding programs could facilitate the selection of resilient cultivars. Field validation should consider both biomass production and forage nutritional quality.

**Highlight:** Time-resolved chlorophyll fluorescence dynamics, combined with leaf *Na*^+^/*K*^+^ homeostasis and carbohydrate partitioning, provide physiological candidate indicators for early salinity tolerance screening in alfalfa.

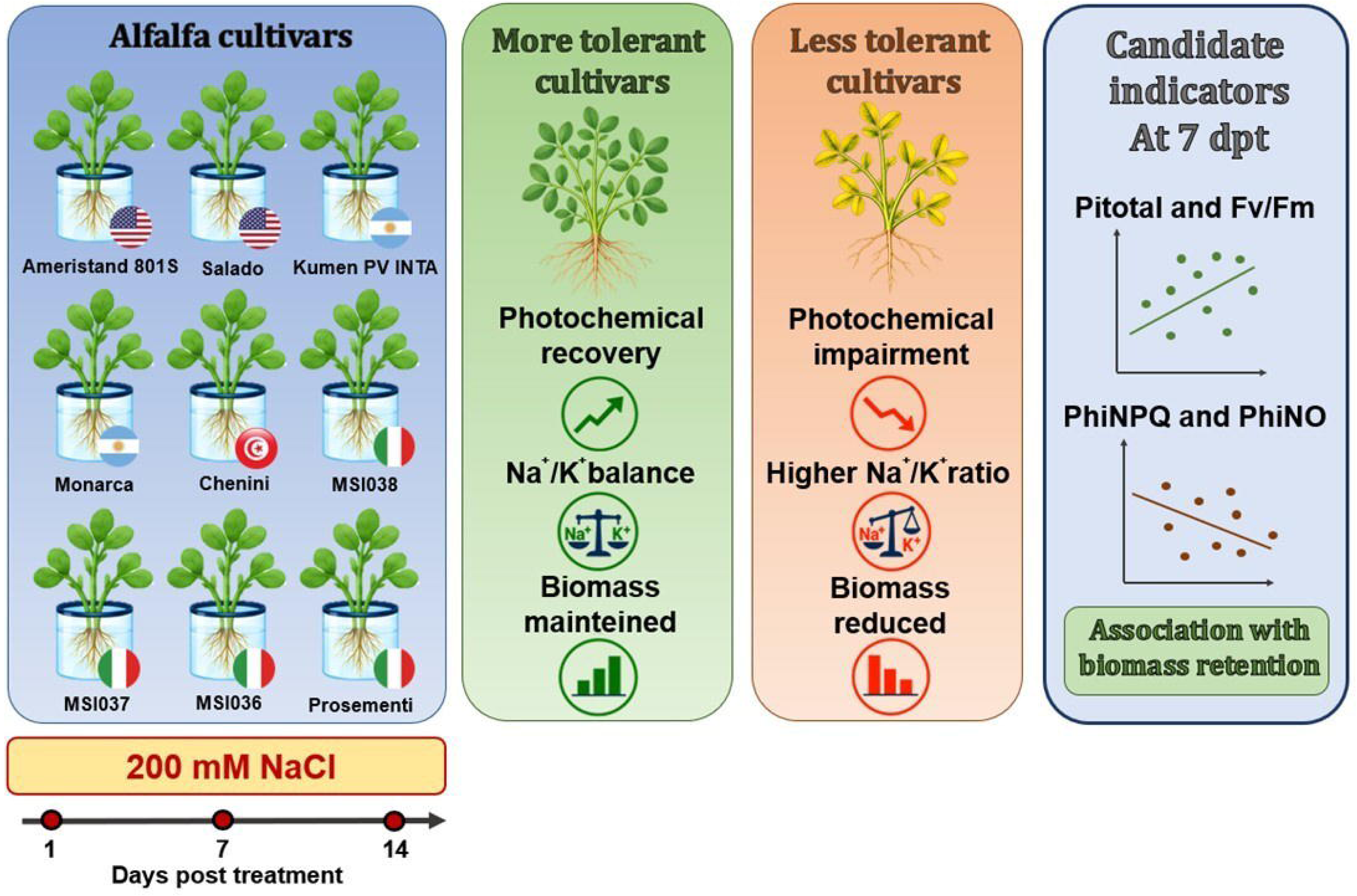

## 1. Introduction

Salinity is one of the major abiotic limitations on agricultural productivity, particularly in irrigated and semi-arid regions where salt accumulation is intensified by limited rainfall, freshwater scarcity and irrigation practices (Munns & Millar 2023). The most recent global assessment by FAO estimates that salt-affected soils cover approximately 1.38 billion ha, representing 10.7% of the global land area, with around 10% of both irrigated and rainfed cropland affected (FAO, 2024). Salinity causes both osmotic and ionic stresses. The early osmotic phase restricts water uptake and carbon assimilation, whereas prolonged exposure promotes Na□ accumulation, nutrient imbalance and oxidative stress, ultimately reducing plant growth and yield (van Zelm *et al*. 2020; Ma *et al*. 2026). Because these effects develop over time, salinity tolerance depends not only on the initial stress response but also on the capacity to re-establish physiological homeostasis during acclimation.

Alfalfa (*Medicago sativa L.*) is the most widely grown perennial legume worldwide (Annicchiarico *et al*. 2015) and represents a cornerstone of global livestock systems, owing to its superior biomass production and nutritional profile. Its cultivation is increasingly constrained by soil salinity and the reliance on marginal irrigation water. Salinity tolerance is already a key factor, along with drought tolerance, for cultivar targeting in various regions, such as the Mediterranean (Annicchiarico et al., 2011). Alfalfa exhibits moderate, cultivar-dependent salt tolerance, offering avenues for genetic improvement (Zhang *et al*. 2024; Pecetti *et al*. 2024). However, the physiological mechanisms coordinating ion homeostasis, carbon metabolism, and growth among alfalfa cultivars under salinity remain incompletely understood. Moreover, while the deleterious effects of salinity on biomass production are well established, its effects on forage quality, including crude protein (CP), fiber fractions (NDF, ADF, ADL), and non-structural carbohydrates (NSC), remain comparatively less documented. In general, forage quality is highly variable and deeply influenced by stress severity and genetic background (Moore & Hatfield 1994; Van Soest 1994). Salinity-induced changes in forage quality are driven by carbon reallocation toward soluble sugar accumulation for osmotic adjustment, thereby altering NSC composition (Sandhu *et al*. 2017). Salinity may influence forage quality through multiple mechanisms, including osmotic effects on plant growth and developmental progression, which can alter fiber accumulation and leaf-to-stem allocation. Similar responses have been documented in drought-stressed alfalfa, where changes in growth and plant morphology were associated with shifts in forage composition (Halim *et al*., 1989). In particular, negative associations between biomass production and forage quality have been reported across diverse alfalfa accessions under water deficit (Zhang *et al*., 2018). Drought-induced increases in leaf-to-stem ratio, digestibility, and crude protein content have also been documented (Carter & Sheaffer, 1983; Pecetti *et al*., 2017). Although the underlying mechanisms and magnitude of these responses may differ between water deficit and salinity, these findings suggest that stress-induced changes in plant growth and biomass partitioning can have important consequences for forage nutritional value. Therefore, assessing salinity tolerance in alfalfa requires considering forage quality alongside biomass production.

Ion homeostasis is a major component of salt tolerance. Under saline conditions, excessive Na□ accumulation in leaves disrupts enzymatic activity, membrane function and nutrient balance, particularly by interfering with K□-dependent metabolic processes (Munns & Millar 2023). Tolerant cultivars often restrict Na□ transport to aerial tissues or maintain a more favorable Na^+^/K^+^ ratio, thereby reducing ionic toxicity (Smethurst *et al*. 2008; Cornacchione & Suarez 2015). However, ion regulation alone does not fully explain differences in cultivar performance, because growth under salinity also depends on the maintenance of photosynthetic activity and carbon supply. Therefore, carbon metabolism adds another layer to salinity acclimation. For instance, soluble sugars can contribute to osmotic adjustment, cellular protection, redox regulation and stress signaling, whereas starch metabolism reflects changes in carbon storage and mobilization between source and sink tissues (Saddhe *et al*. 2021; Ribeiro *et al*. 2022). Under salinity, sustained carbon supply is essential for root maintenance, ion transport, and osmotic regulation. Thus, carbohydrate dynamics may help explain why some cultivars recover photosynthetic function and maintain growth, whereas others experience carbon limitation and growth inhibition.

Photosynthesis is one of the earliest physiological processes affected by salinity (Chaves *et al*. 2009). Reduced CO□ assimilation increases excitation pressure within the photosynthetic apparatus, leading to lower photosystem II (PSII) efficiency, altered electron transport and activation of energy dissipation mechanisms that protect against photodamage (Baker 2008). Chlorophyll a fluorescence (ChlF) therefore provides a sensitive approach to detect stress-induced changes in photosynthetic function before visible damage becomes evident (Murchie & Lawson 2013). Measurements over time can distinguish transient photoprotective adjustment from persistent photoinhibition. This distinction is particularly relevant for cultivar screening, because tolerance may depend more on photosynthetic stability than on the magnitude of the initial decline.

Developing alfalfa cultivars with enhanced salt tolerance and forage quality is critical for sustaining agricultural production in increasingly saline environments. In this study, we evaluated nine alfalfa cultivars previously characterized and selected for their diverse responses to salinity (Pecetti et al., 2024). We hypothesized that i) photosynthetic acclimation following an acute salt shock, rather than initial baseline resistance, determines final biomass preservation across cultivars; and ii) salinity decouple from overall forage nutritional value due to non-structural carbohydrate alteration and cell wall lignification. Specifically, we investigated whether cultivars that maintain biomass under salt stress also maintain PSII efficiency following an initial decline, and whether this stability correlates with distinct leaf Na^+^/K^+^ ratios or carbohydrate partitioning. A secondary objective was to determine the impact of salinity on key forage quality traits. To achieve these aims, we analyzed biometric parameters, leaf ion homeostasis (Na□and K□), shoot and root soluble sugar and starch profiles, and chlorophyll fluorescence (ChlF) dynamics. By integrating these physiological dimensions, we aimed to identify physiological and photosynthetic indicators integrated into predictive multivariate models for salt tolerance in alfalfa, ultimately contributing to the selection of more resilient cultivars. To this end, we applied an acute 200 mM NaCl treatment under hydroponic conditions to elicit contrasting physiological responses among cultivars and facilitate the identification of discriminative traits. This approach provides a framework for rapid phenotypic screening that could support alfalfa breeding programs targeting saline environments.

## 2. Materials and Methods

### 2.1. Plant material and experimental setup

Seeds of nine alfalfa (*Medicago sativa* L.) cultivars were used, including Ameristand 801S and Salado bred in the USA, the Argentinian cultivars Kumen and Monarca, the Italian cultivar Prosementi, the Tunisian landrace Chenini, and CREA’s selections MSI036, MSI037 and MSI038. All cultivars except Prosementi underwent a previous field evaluation in either or both Northern Argentina under saline soil and Tunisia under irrigation with saline water (Pecetti et al., 2024), showing salt tolerance ranging from fairly modest to high partly as a function of the test region. Prosementi represented a salt less tolerant reference cultivar based on results in Annicchiarico *et al*. (2011) and complementary results in Pecetti *et al*. (2024).

Seeds of each cultivar were germinated on mesh supports containing moist vermiculite positioned over 3-L containers filled with five-fold diluted B&D nutrient solution (Broughton & Dilworth, 1971), supplemented with NH NO to a final concentration of 5 mM under continuous aeration. The nutrient solution was adjusted to pH 6.5 and renewed weekly. Plants were grown in a controlled-environment growth chamber under a 16 h light/8 h dark photoperiod, a photosynthetic photon flux density of 300 μmol m−² s −¹, and a temperature of 23 ± 2 °C.

Three independent experiments were conducted. Salt stress was imposed 20 d after sowing by adding a shock of NaCl 200 mM. The acute 200 mM NaCl shock was selected based on preliminary screening assays establishing this concentration as the optimal threshold to maximize inter-cultivar physiological variance and induce rapid metabolic reprogramming without causing unspecific osmotic collapse within 14 days. Control plants remained in nutrient solution without NaCl. Leaf and root samples were collected 14 days post treatment (dpt). Samples or biochemical analyses were immediately frozen in liquid nitrogen and stored at −80 °C until further analysis (Fig. S1).

### 2.2. Biometric parameters

Biometric measurements were recorded for both control and salt-treated plants at the end of the experimental period, based on three independent experiments with thirty plants per treatment (n = 3). The following variables were assessed: shoot length (cm), main root length (cm), and total leaf number per plant. For biomass determination, shoots and roots were separately harvested and oven-dried at 40 °C for 72 h until constant weight.

### 2.3. Forage quality parameters

Forage nutritional quality was assessed using near-infrared reflectance spectroscopy (NIRS, DS2500 FOSS). Samples were obtained from three independent experiments, each comprising 30 plants per treatment pooled into a single sample (n = 3). Ground samples (1-mm particle size) were scanned in reflectance mode over the 400–2500 nm wavelength range at 2-nm intervals, with 32 co-added scans per spectrum. Calibration equations were developed using reference samples for which crude protein (CP; n = 861), ash (n = 424), neutral detergent fiber (NDF; n = 594), acid detergent fiber (ADF; n = 1035), and acid detergent lignin (ADL; n = 167) concentrations had been determined by wet-chemistry methods according to AOAC (1990). Spectral data were pre-processed using standard normal variate (SNV) and detrending transformations, and calibration equations were developed by partial least squares (PLS) regression using leave-one-out cross-validation. All experimental samples were analyzed in duplicate, and predicted values were expressed as a percentage of dry matter (% DM).

### 2.4. Ion concentration

Leaf Na□ and K□ were extracted from dried leaf powder in 30% (v/v) methanol and quantified by non-suppressed ion chromatography with conductivity detection (Shim-pack IC-C3 column; 3 mM oxalic acid as the mobile phase; 40 °C; 1.2 mL min ¹), following Pantsar-Kallio & Manninen (1995). Na□ and K□ were determined by HPLC (Shimadzu Prominence Modular HPLC, Kyoto, Japan). Chromatographic runs were performed at 40 °C with a flow rate of 1.2 mL min ¹ for 18 min under non-suppressed conditions, and ions were detected by conductivity. Chromatography was registered and analyzed using LabSolutions (ver. 5.6) software (Pantsar-Kallio & Manninen 1995). Three plants per treatment were pooled in each of three independent experiments (n = 3).

### 2.5. Soluble Sugars (SS) content

Total soluble sugars (SS) were quantified by the anthrone method (Fales 1951), adapted to a final reaction volume of 200 µL. Ethanolic leaf and root extract was mixed with anthrone reagent, heated at 100 °C for 5 min, rapidly cooled on ice, and absorbance was measured at 620 nm. Glucose was used for calibration. Three plants per treatment were pooled in each of three independent experiments (n = 3).

### 2.6. Starch content

Starch content was determined in leaf and root pellets by quantifying the reducing sugars released following hydrolysis of the pellet with α-amyloglucosidase (Sumner & Somers 1944), using glucose as the standard. Values were expressed as mg glucose equivalents g ¹ DW. Three plants per treatment were pooled in each of three independent experiments (n = 3).

### 2.7. Chlorophyll-*a* fluorescence and photosynthetic performance measurements

ChlF and photosynthetic performance measurements were conducted at 1, 7, and 14 dpt. Fast ChlF transients were recorded using a Pocket-PEA fluorometer (Plant Efficiency Analyzer; Hansatech Instruments Ltd., King’s Lynn, Norfolk, UK). The third fully expanded leaf of 12 plants per cultivar and treatment in each of three independent experiments (36 plants per cultivar and treatment in total) were dark-adapted for at least 30 min using leaf clips to ensure complete oxidation of the PSII reaction centers. Leaves were subsequently exposed to a saturating light pulse of 3500 μmol photons m−² s −¹ for 1 s. The OJIP transients were analyzed according to the JIP-test (Strasser et al., 2004). Derived parameters included the absorption flux per active PSII reaction center (ABS/RC), electron transport flux beyond Q□per active PSII reaction center (ET□/RC), energy dissipation flux per active PSII reaction center (DI□/RC), electron flux reducing terminal electron acceptors at the PSI acceptor side per active PSII reaction center (RE□/RC), and the total performance index (PI□□□□□). Definitions of all calculated parameters are provided in Table S1.

Complementary measurements were conducted at midday on the third fully expanded leaves using a MultispeQ V2.0 handheld device connected to the PhotosynQ platform (Kuhlgert *et al*. 2016), following the publicly available “Photosynthesis Rides 500” protocol. The parameters obtained included the maximum efficiency of PSII in the light-adapted state (Fv’/Fm’), effective quantum yield of PSII photochemistry (Phi2), estimated linear electron flow (LEF), the fraction of open PSII reaction centers (qL), proton conductivity of the thylakoid membrane (gH□), steady-state proton flux across the thylakoid membrane (vH□), non-photochemical quenching (NPQt), and the quantum yields of regulated (PhiNPQ) and non-regulated (PhiNO) energy dissipation (Table S1).

### 2.8. Statistical Analysis

All experiments were performed with at least three biological replicates. Statistical significance among means was assessed by analysis of variance (ANOVA), followed by Tukey’s post hoc test (*P* < 0.05). Data processing and statistical analyses were conducted using InfoStat software (Di Rienzo *et al.,* 2019). Spearman correlation analyses were performed using GraphPad Prism version 10.0.0 for macOS (GraphPad Software, Boston, MA, USA). Percentage changes relative to the control were calculated as [(treatment mean − control mean)/control mean] × 100, with positive and negative values indicating increases and decreases, respectively.

For the integrative analyses, data were summarized at the cultivar level (n = 9). Shoot dry weight values from the six biological replicates available for each cultivar and treatment were averaged, and biomass retention under salinity was calculated as the ratio between mean shoot dry weight under NaCl treatment and mean shoot dry weight under control conditions (NaCl/Control). This ratio was used as a quantitative indicator of cultivar performance under salinity.

Multivariate analyses were performed in Python using the pandas, SciPy, and scikit-learn libraries. Principal component analysis (PCA) was conducted using the 52 physiological, chlorophyll fluorescence (ChlF), ionic, and biochemical variables measured in the study. For each variable, the salinity response was expressed as a log□fold change (NaCl/Control), and the resulting data matrix was standardized prior to PCA. The association between PC1 scores and biomass retention was assessed using Spearman’s rank correlation.

Pairwise Spearman correlations were also calculated among the 52 measured variables and biomass retention across the nine cultivars, resulting in 1,378 unique pairwise comparisons. P-values were adjusted for multiple testing using the Benjamini–Hochberg false discovery rate (FDR) procedure, with statistical significance defined as an FDR-adjusted P < 0.05.

## 3. Results

### 3.1. Salinity induces contrasting shoot growth preservation and root biomass allocation across alfalfa cultivars

The aerial response to 200 mM NaCl varied among the nine evaluated cultivars (Table 1, Fig. S2). Ameristand 801S, Salado, MSI038, Kumen, and Chenini exhibited better ability to maintain shoot growth under stress relative to control conditions, with small, nonsignificant reductions in the measured traits (P > 0.05). Consistent with these responses, shoot biomass retention under salinity was highest in Kumen and Ameristand 801S (Fig. 1). In contrast, Prosementi exhibited the greatest overall susceptibility to salinity, suffering high and significant (P < 0.05) reductions in shoot DW (-32.0%), leaf number and shoot length relative to the control. Monarca and MSI037 exhibited a significant decrease in all growth traits, whereas MSI036 showed a decrease in shoot DW combined with a relatively stable shoot length.

**Fig. 1.**
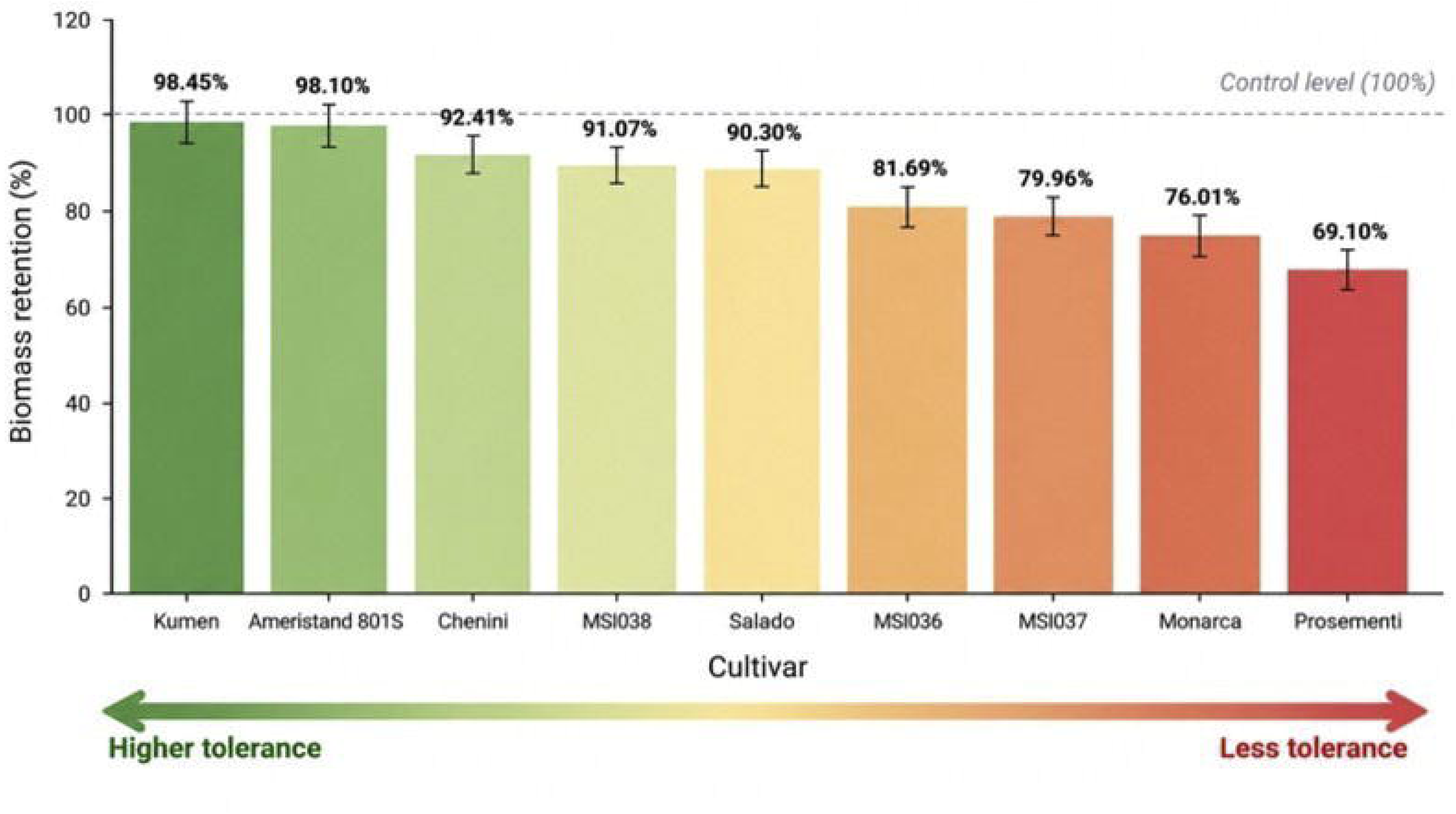
Biomass retention of alfalfa cultivars under salinity stress. Biomass retention was calculated as the ratio between shoot dry weight under 200 mM NaCl and the corresponding control condition (NaCl/Control), expressed as a percentage. Cultivars are ranked from highest to lowest biomass retention. The dashed horizontal line indicates the control level (100%), representing no reduction in shoot biomass under salinity. Higher biomass retention indicates a greater ability to maintain biomass production under salt stress. The gradient from green to red represents decreasing tolerance to salinity. Values were calculated from three biological replicates per cultivar and treatment.

**Table 1.**
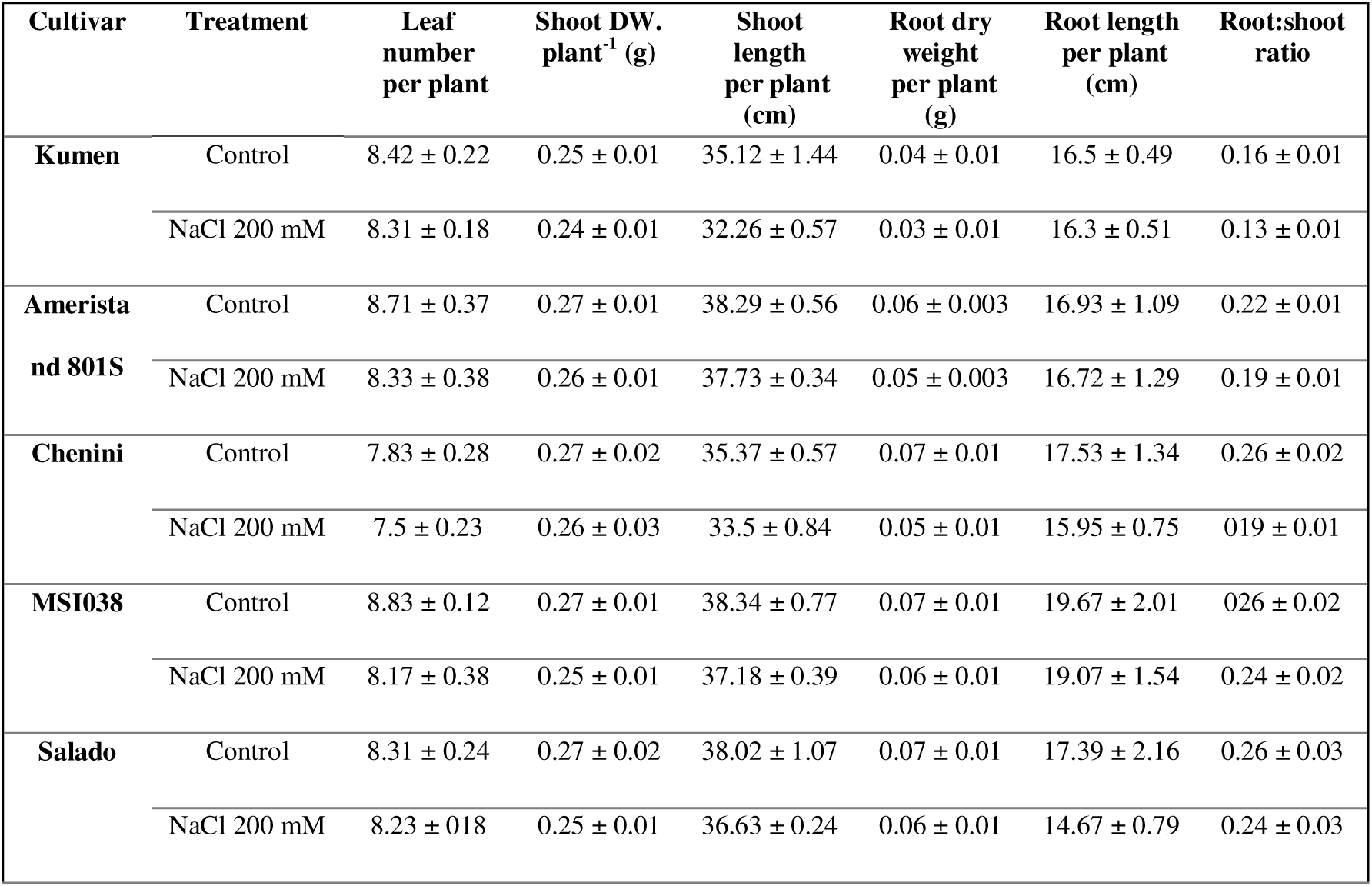
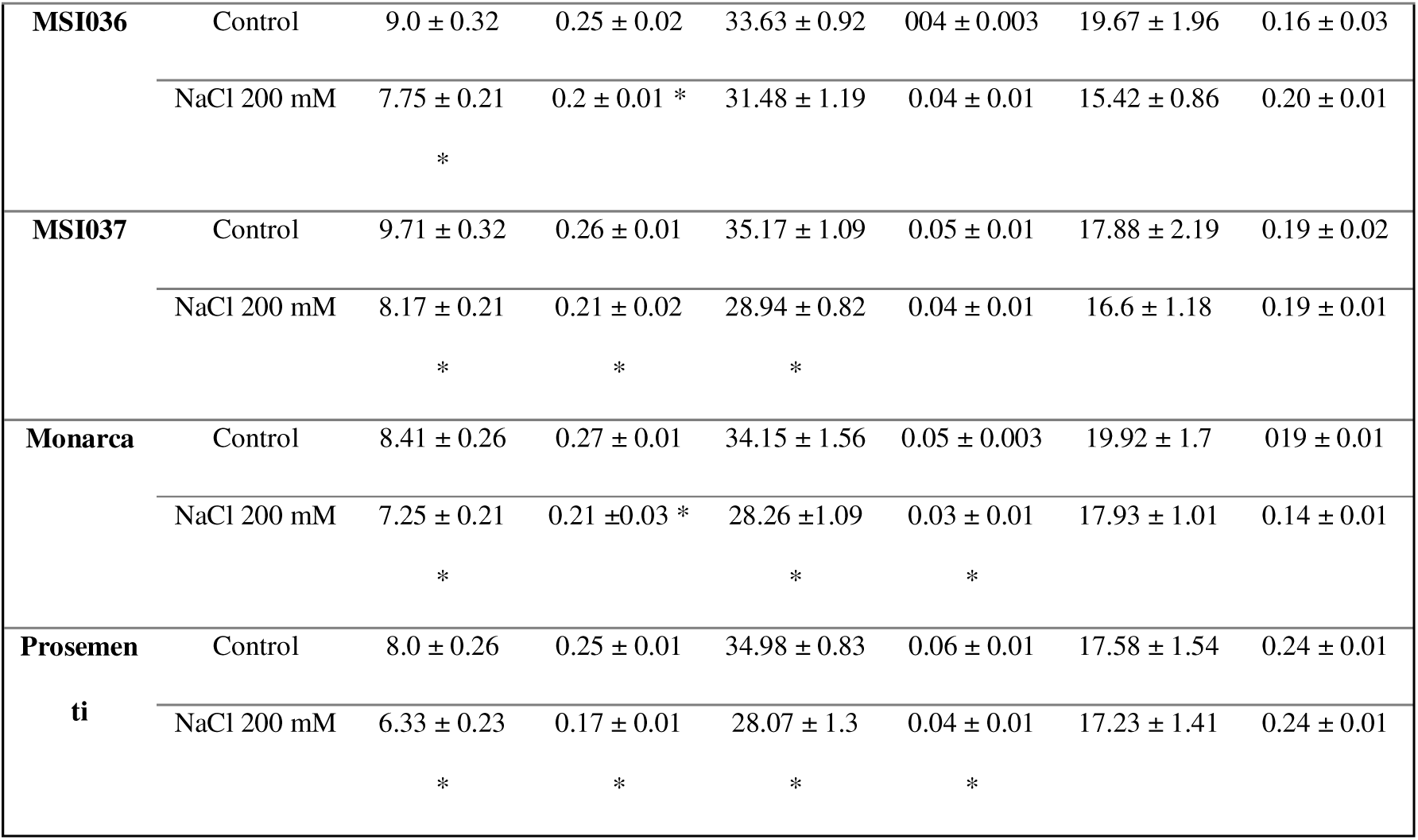
Effect of salt stress on growth traits of alfalfa cultivars. Leaf number, shoot dry weight, shoot length, root dry weight, root length and root:shoot ratio measured in nine alfalfa cultivars grown under control conditions and exposed to 200 mM NaCl. Values represent means ± SE. Asterisks indicate significant differences between plants under saline treatment and their respective controls, based on three independent experiments with thirty plants per treatment (n=3), analyzed using ANOVA followed by Tukey’s test (p < 0.05).

| **Cultivar** | **Treatment** | **Leaf number per plant** | **Shoot DW. plant<sup>-1</sup> (g)** | **Shoot length per plant (cm)** | **Root dry weight per plant (g)** | **Root length per plant (cm)** | **Root:shoot ratio** |
|---|---|---|---|---|---|---|---|
| **Kumen** | Control | 8.42 ± 0.22 | 0.25 ± 0.01 | 35.12 ± 1.44 | 0.04 ± 0.01 | 16.5 ± 0.49 | 0.16 ± 0.01 |
|  | NaCl 200 mM | 8.31 ± 0.18 | 0.24 ± 0.01 | 32.26 ± 0.57 | 0.03 ± 0.01 | 16.3 ± 0.51 | 0.13 ± 0.01 |
| **Ameristand 801S** | Control | 8.71 ± 0.37 | 0.27 ± 0.01 | 38.29 ± 0.56 | 0.06 ± 0.003 | 16.93 ± 1.09 | 0.22 ± 0.01 |
|  | NaCl 200 mM | 8.33 ± 0.38 | 0.26 ± 0.01 | 37.73 ± 0.34 | 0.05 ± 0.003 | 16.72 ± 1.29 | 0.19 ± 0.01 |
| **Chenini** | Control | 7.83 ± 0.28 | 0.27 ± 0.02 | 35.37 ± 0.57 | 0.07 ± 0.01 | 17.53 ± 1.34 | 0.26 ± 0.02 |
|  | NaCl 200 mM | 7.5 ± 0.23 | 0.26 ± 0.03 | 33.5 ± 0.84 | 0.05 ± 0.01 | 15.95 ± 0.75 | 019 ± 0.01 |
| **MSI038** | Control | 8.83 ± 0.12 | 0.27 ± 0.01 | 38.34 ± 0.77 | 0.07 ± 0.01 | 19.67 ± 2.01 | 026 ± 0.02 |
|  | NaCl 200 mM | 8.17 ± 0.38 | 0.25 ± 0.01 | 37.18 ± 0.39 | 0.06 ± 0.01 | 19.07 ± 1.54 | 0.24 ± 0.02 |
| **Salado** | Control | 8.31 ± 0.24 | 0.27 ± 0.02 | 38.02 ± 1.07 | 0.07 ± 0.01 | 17.39 ± 2.16 | 0.26 ± 0.03 |
|  | NaCl 200 mM | 8.23 ± 018 | 0.25 ± 0.01 | 36.63 ± 0.24 | 0.06 ± 0.01 | 14.67 ± 0.79 | 0.24 ± 0.03 |
| **MSI036** | Control | 9.0 ± 0.32 | 0.25 ± 0.02 | 33.63 ± 0.92 | 004 ± 0.003 | 19.67 ± 1.96 | 0.16 ± 0.03 |
|  | NaCl 200 mM | 7.75 ± 0.21 * | 0.2 ± 0.01 * | 31.48 ± 1.19 | 0.04 ± 0.01 | 15.42 ± 0.86 | 0.20 ± 0.01 |
| **MSI037** | Control | 9.71 ± 0.32 | 0.26 ± 0.01 | 35.17 ± 1.09 | 0.05 ± 0.01 | 17.88 ± 2.19 | 0.19 ± 0.02 |
|  | NaCl 200 mM | 8.17 ± 0.21 * | 0.21 ± 0.02 * | 28.94 ± 0.82 * | 0.04 ± 0.01 | 16.6 ± 1.18 | 0.19 ± 0.01 |
| **Monarca** | Control | 8.41 ± 0.26 | 0.27 ± 0.01 | 34.15 ± 1.56 | 0.05 ± 0.003 | 19.92 ± 1.7 | 019 ± 0.01 |
|  | NaCl 200 mM | 7.25 ± 0.21 * | 0.21 ±0.03 * | 28.26 ±1.09 * | 0.03 ± 0.01 * | 17.93 ± 1.01 | 0.14 ± 0.01 |
| **Prosementi** | Control | 8.0 ± 0.26 | 0.25 ± 0.01 | 34.98 ± 0.83 | 0.06 ± 0.01 | 17.58 ± 1.54 | 0.24 ± 0.01 |
|  | NaCl 200 mM | 6.33 ± 0.23 * | 0.17 ± 0.01 * | 28.07 ± 1.3 * | 0.04 ± 0.01 * | 17.23 ± 1.41 | 0.24 ± 0.01 |

The analysis of the root traits revealed cultivar responses that were partly different from those for shoot traits (Table 1). The accumulation of root biomass was highly sensitive in specific cultivars, yet longitudinal root elongation remained unaffected across the board. Root DW was significantly reduced in Monarca and Prosementi by 40.0% and 33.3%, respectively, relative to their corresponding cultivar controls (P < 0.05). In contrast, no statistically significant reductions in root length were detected for any of the other cultivars. These dynamics indicate that the reduced root biomass accumulation observed in less tolerant cultivars occurred without a detectable inhibition of primary root elongation.

### 3.2. Salinity Alters Alfalfa Forage Quality through Cultivar-Specific Shifts in Protein, Structural Fiber, and Non-Structural Carbohydrates

NIRS-based estimates revealed shifts in forage quality, with significant cultivar-dependent responses to salt stress (Table 2). For example, total Ash% content declined significantly (P < 0.05) across all cultivars, with reductions ranging from -16.6% in Ameristand 801S to -28.6% in Monarca. CP content exhibited significant stress-induced increases (P < 0.05) in six cultivars and especially in MSI038 (+50.3%), Salado (+48.3%), MSI037 (+43.6%), and MSI036 (+40.8%). CP concentration did not differ significantly from the corresponding control in Ameristand 801S, Prosementi, and Monarca. The structural carbohydrate matrix, which dictates forage digestibility and voluntary intake limits, demonstrated complex, divergent responses among cultivars. ADL increased significantly in MSI037 (+19.5%), MSI038 (+16.3%), Ameristand 801S (+11.3%), and MSI036 (+11.0%). In contrast, ADF decreased significantly in Monarca (−12.1%), Prosementi (−10.4%), and Ameristand 801S (−9.4%). NDF decreased significantly in Ameristand 801S (−6.2%) but increased significantly in MSI036 (+8.2%) and MSI038 (+3.8%). Non-structural carbohydrate (NSC) content, calculated as the sum of independently measured soluble sugars and starch, was lower under salinity in all cultivars, with decreases ranging from 6.4% in Chenini to 57.2% in MSI036. The largest reductions occurred in MSI036 (−57.2%), Monarca (−48.7%), and Prosementi (−43.6%).

**Table 2.** Salinity-induced shifts in forage nutritive value among alfalfa cultivars. Crude protein (CP), ash, acid detergent lignin (ADL), acid detergent fiber (ADF), and neutral detergent fiber (NDF) were estimated by near-infrared reflectance spectroscopy (NIRS) in nine alfalfa cultivars grown under control conditions or exposed to 200 mM NaCl. Non-structural carbohydrates (NSC) were calculated as the sum of independently measured soluble sugar and starch contents. Values represent means ± SE from three independent experiments, each comprising 30 plants per treatment pooled into a single sample (n = 3). Asterisks indicate significant differences between NaCl-treated plants and their respective controls, as determined by ANOVA followed by Tukey’s test (*P* < 0.05).

| Cultivar | Treatment | CP% | Ash% | ADL% | ADF% | NDF% | NSC% |
|---|---|---|---|---|---|---|---|
| Kumen | Control | 15.93 ± 0.81 | 6.49 ± 0.12 | 9.13 ± 0.26 | 34.55 ± 0.41 | 48.09 ± 0.88 | 0.42 ± 0.03 |
|  | NaCl 200 mM | 20.4 ± 0.34 * | 4.98 ± 0.09 * | 9.75 ± 0.16 | 33.07 ± 0.3 * | 50.33 ± 0.78 | 0.30 ± 0.03 |
| Ameristand 801S | Control | 16.86 ± 0.99 | 6.52 ± 0.1 | 8.75 ± 0.17 | 36.18 ± 0.27 | 50.86 ± 0.73 | 0.34 ± 0.03 |
|  | NaCl 200 mM | 17.52 ± 0.56 | 5.44 ± 0.21 * | 9.74 ± 0.18 * | 32.77 ± 0.44 * | 47.7 ± 0.39 * | 0.32 ± 0.04 |
| Chenini | Control | 15.34 ± 0.7 | 6.2 ± 0.11 | 9.14 ± 0.25 | 33.63 ± 1.01 | 47.5 ± 2.17 | 0.37 ± 0.02 |
|  | NaCl 200 mM | 18.8 ± 0.99 * | 4.79 ± 0.09 * | 9.69 ± 0.2 | 33.31 ± 0.43 | 51.56 ± 0.38 | 0.35 ± 0.03 |
| MSI038 | Control | 14.62 ± 0.76 | 5.7 ± 0.17 | 9.09 ± 0.14 | 30.68 ± 0.55 | 45.69 ± 0.42 | 0.45 ± 0.06 |
|  | NaCl 200 mM | 21.98 ± 1.14 * | 4.1 ± 0.16 * | 10.57 ± 0.05 * | 30.03 ± 0.65 | 47.44 ± 0.46 * | 0.39 ± 0.05 |
| Salado | Control | 13.82 ± 1.1 | 6.37 ± 0.47 | 9.27 ± 0.51 | 33.87 ± 1.67 | 47.84 ± 1.04 | 0.35 ± 0.03 |
|  | NaCl 200 mM | 20.5 ± 2.42 * | 4.84 ± 0.27 * | 10.79 ± 0.6 | 32.22 ± 1.02 | 48.57 ± 1.59 | 0.28 ± 0.01 |
| MSI036 | Control | 15.26 ± 0.76 | 5.62 ± 0.07 | 9.47 ± 0.11 | 31.05 ± 0.27 | 45.01 ± 0.32 | 0.40 ± 0.05 |
|  | NaCl 200 mM | 21.48 ± 0.8 * | 4.39 ± 0.11 * | 10.51 ± 0.43 * | 31.25 ± 0.75 | 48.7 ± 0.99 * | 0.17 ± 0.03 * |
| MSI037 | Control | 13.48 ± 0.19 | 6.22 ± 0.1 | 8.82 ± 0.16 | 32.83 ± 0.8 | 46.9 ± 1.27 | 0.34 ± 0.02 |
|  | NaCl 200 mM | 19.36 ± 1.8 * | 4.57 ± 0.07 * | 10.54 ± 0.18 * | 31.41 ± 0.25 | 48.12 ± 0.45 | 0.25 ± 0.01 * |
| Monarca | Control | 17.63 ± 0.56 | 6.36 ± 0.12 | 10.06 ± 0.09 | 35.48 ± 0.87 | 49.16 ± 1.32 | 0.51 ± 0.03 |
|  | NaCl 200 mM | 18.02 ± 0.23 | 4.54 ± 0.11 * | 10.36 ± 0.3 | 31.19 ± 0.2 * | 47.69 ± 0.27 | 0.26 ± 0.03 * |
| Prosementi | Control | 15.72 ± 2.31 | 6.99 ± 0.15 | 8.89 ± 0.49 | 37.75 ± 0.56 | 51.81 ± 0.85 | 0.47 ± 0.03 |
|  | NaCl 200 mM | 17.42 ± 0.92 | 5.09 ± 0.16 * | 9.46 ± 0.28 | 33.82 ± 0.51 * | 52.68 ± 0.64 | 0.27 ± 0.04 * |

### 3.3. Disruption of Leaf Ion Homeostasis and Na□ Accumulation Are Associated with Less Tolerant Cultivars

Leaf Na□ concentration increased significantly in all alfalfa cultivars under salt stress compared with their respective control (Fig. 2A). However, the extent of this ion accumulation revealed statistically significant cultivar variations. The more salt-tolerant group based on the biomass results (Chenini, Ameristand 801S, Kumen, and MSI038) showed comparatively lower foliar Na□ accumulation. In contrast, Prosementi, Monarca, and MSI036 accumulated substantially higher foliar Na□ concentrations. Several cultivars also exhibited significant increases in K□ concentration under salinity (P < 0.05) (Fig. 2B). While Ameristand 801S, MSI038, and Chenini showed no significant changes in K□ concentration; Salado, Kumen, MSI036, Monarca, and Prosementi showed significant increases. The cultivar MSI037 showed a distinct ionic response, combining high Na□ accumulation with a marked and significant increase in K□ concentration. Some cultivars showed comparatively smaller increases in the Na^+^/K^+^ ratio under salinity (Fig. 2C). This group included Kumen, Chenini, Ameristand 801S, and MSI037. Cultivars MSI038 and Salado occupied intermediate positions. Na^+^/K^+^ under NaCl was highest in MSI036, Monarca and Prosementi (Fig. 2C; Table S2).

**Fig. 2.**
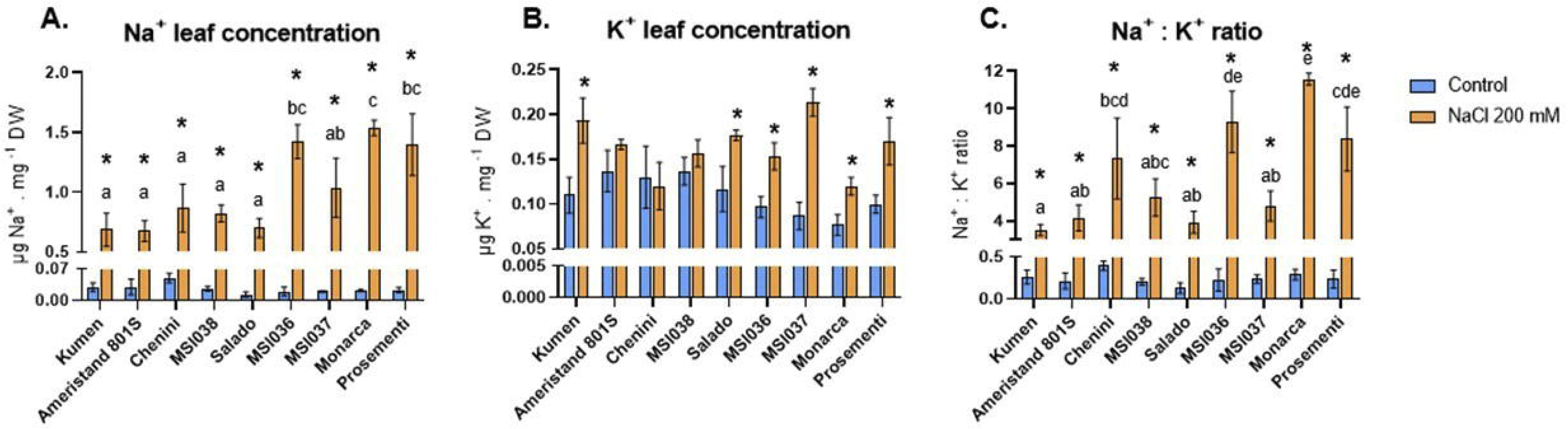
Effect of salinity on leaf ion concentrations in alfalfa cultivars. (A) Sodium (Na□) concentration, (B) potassium (K□) concentration, and (C) Na□:K□ ratio in leaves of nine alfalfa cultivars under control conditions and 200 mM NaCl. Data are presented as means ± SE from three independent biological experiments, with each biological replicate consisting of a pool of three plants. Asterisks indicate significant differences between control and NaCl treatments within each cultivar, whereas different lowercase letters indicate significant differences among cultivars under NaCl treatment. Statistical differences were determined by two-way ANOVA with treatment and cultivar as factors, followed by Tukey’s post-hoc test (*P* < 0.05).

### 3.4. Salinity Elicits Cultivar-Specific Reprogramming of Source–Sink Carbohydrate Partitioning and Soluble Sugar-to-Starch Ratios

Whole-plant carbohydrate dynamics (Table S3) provided insights into the relationship between carbon assimilation, carbohydrate allocation, and biomass accumulation, helping interpret the observed phenotypic responses. Salt stress induced cultivar-specific shifts in carbon homeostasis, as evidenced by changes in soluble sugar and starch pools, reflecting adjustments in readily available carbohydrates involved in transport and signaling and in transient carbon and energy reserves, respectively. Within the foliar source tissues, soluble sugar concentrations exhibited divergent, statistically significant responses (P < 0.05) to salinity. A marked accumulation of soluble sugars was observed in MSI037 (+79.7%), MSI038 (+46.6%), and Chenini (+38.7%). In contrast, Monarca, Kumen, Prosementi, and MSI036 showed decrease in their foliar soluble sugar, ranging from -43.6 to -24.2%. Foliar starch content was severely and significantly (P < 0.05) decreased in MSI036 (-63.8%), Monarca (-49.4%), Prosementi (-46.7%), and MSI037 (-45.9%). This collective metabolic adjustment altered the foliar soluble sugar-to-starch ratio (Soluble sugar/Starch) (Fig. 3A), with a particularly high increase of the ratio observed in MSI037 (+232.31%), MSI036 (+109.60%), and MSI038 (+96.11%). The increased soluble sugar/starch ratio was consistent with a stress-induced shift in carbohydrate partitioning toward soluble sugars relative to starch, potentially contributing to osmotic adjustment.

**Fig. 3.**
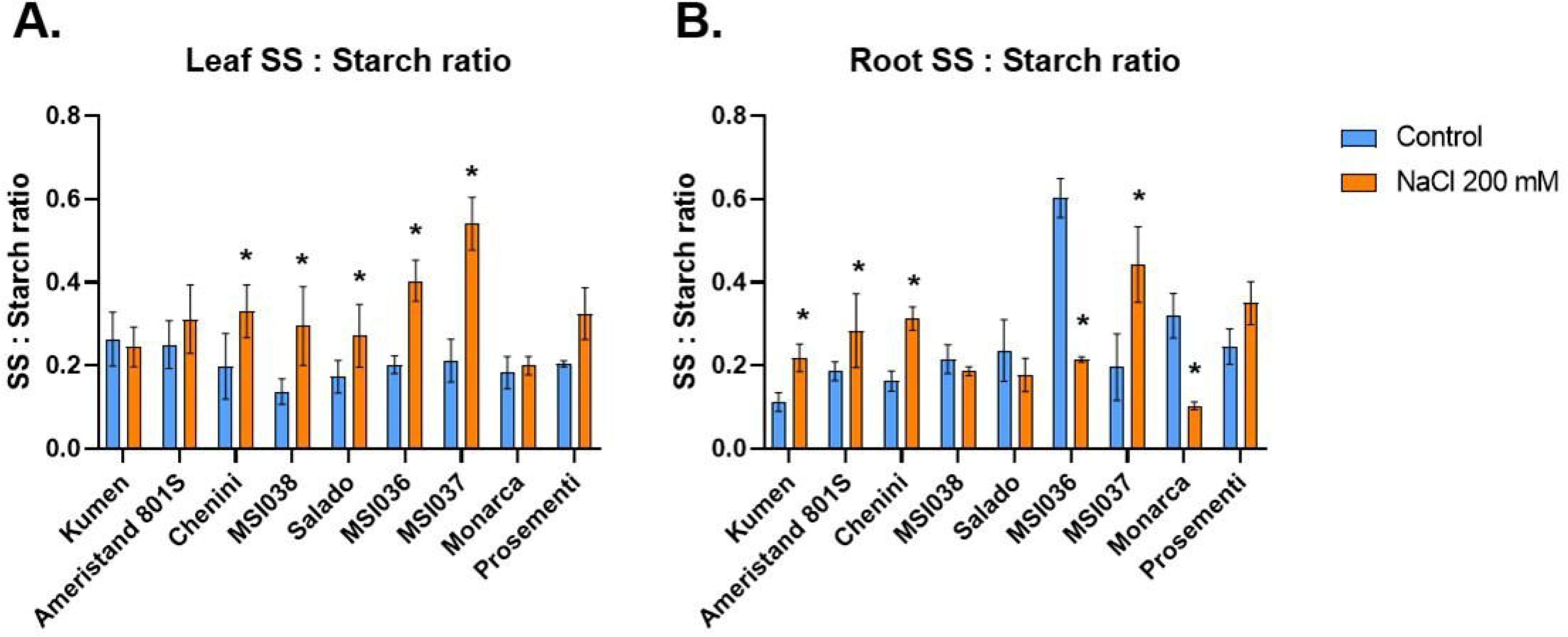
Organ-specific carbohydrate adjustments in alfalfa cultivars under salinity. Soluble sugar-to-starch ratio in (A) leaves and (B) roots of nine alfalfa cultivars grown under control conditions or exposed to 200 mM NaCl for 14 days. Values represent means ± SE. Asterisks indicate significant differences between plants under saline treatment and their respective controls, based on three independent experiments with three plants per treatment, grouped into one pool (n=3), analyzed using ANOVA followed by Tukey’s test (p < 0.05).

The partitioning of carbohydrates to the root system, the primary sink during osmotic stress, further outlined the different cultivar adaptive strategies for stress tolerance. Soluble sugar content in roots increased significantly (P < 0.05) in Prosementi (+77.8%), MSI037 (+64.3%), Kumen (+25.7%), and Chenini (+22.9%), suggesting an enhanced requirement for active osmolytes or metabolic signaling molecules within the root organ to sustain growth under saline conditions. In contrast, Monarca and MSI036 exhibited a substantial and significant decline in root soluble sugars (-50.0% and -46.6%, respectively). Root starch content also showed cultivar-dependent responses. Salado and Monarca significantly increased root starch by 52.8% and 46.9%, respectively, whereas Kumen, Chenini, Ameristand 801S, and MSI037 showed significant decrease of root starch reserves, ranging from approximately -29.1 to -38.1%. Consequently, the root soluble sugar/starch ratio revealed contrasting carbon allocation patterns across cultivars (Fig. 3B), with MSI037 (+131.8%) and Kumen (+103.3%) showing a shift toward soluble sugars relative to stored non-structural carbohydrate, whereas Monarca (−65.9%) and MSI036 (−63.7%) showed the opposite pattern, characterized by a marked decrease in soluble sugars relative to their starch pools (Table S3). Figure 3B shows that the MSI036 control (ratio ∼0.6) was substantially higher than those of the other cultivars (∼0.1–0.25); therefore, the −63.7% reduction observed in this cultivar is largely influenced by an atypical control value.

### 3.5. Time-resolved ChlF measurements

Salt stress induced distinct cultivar-specific temporal shifts in photochemical efficiency respect to control (Fig. S3). At 1 dpt, genotypic differences were primarily evident in the magnitude and nature of the early photochemical response. Ameristand 801S showed an increase in PhiNPQ together with only minor decreases in ETo/RC, REo/RC, and Phi2, indicating relatively limited early perturbation of photochemical performance. In contrast, MSI038, Salado, Kumen, and Chenini exhibited a broader early response characterized by reductions in Phi2, ETo/RC, REo/RC, and PI_total_, accompanied by rapid increases in NPQt and PhiNPQ, consistent with enhanced regulated energy dissipation. Monarca and MSI037 exhibited immediate photochemical impairment across several parameters, including Fv’/Fm’, Phi2, LEF, and PI_total_, whereas Prosementi displayed no detectable alterations at this initial stage.

By 7 dpt, physiological differences respect to control among cultivars became more apparent. Ameristand 801S maintained marked photochemical stability, exhibiting only localized reductions in REo/RC. MSI038 and Chenini demonstrated partial photochemical stability, with residual impairment largely restricted to electron-transport-related parameters. Salado and Kumen maintained enhanced regulated energy dissipation together with moderate reductions in ETo/RC and LEF. In contrast, Prosementi, Monarca, and MSI037 exhibited persistent photochemical impairment, characterized by reductions in PI_total_, Fv’/Fm’, Phi2, and LEF, together with increases in ABS/RC and DI /RC, sustained regulated energy dissipation (NPQt and PhiNPQ), and increased non-regulated energy dissipation (PhiNO).

By 14 dpt, cultivar-specific differences in photochemical acclimation became more pronounced. Chenini and Kumen showed the strongest preservation of photochemical performance, with values approaching those of their respective controls, whereas Ameristand 801S and MSI038 exhibited only minor residual impairments. Salado displayed an intermediate response, with persistent reductions in ETo/RC and LEF. In contrast, Prosementi and Monarca, and to a lesser extent MSI037 and MSI036, exhibited sustained declines in PSII photochemical efficiency and electron transport, as indicated by reductions in Fv’/Fm’, Phi2, LEF, and REo/RC, together with increased non-photochemical energy dissipation. These alterations were associated with greater biomass penalties under salinity.

Consistent with these endpoint differences, the temporal analysis of Phi2 and PhiNPQ between 1 and 14 dpt revealed distinct photosynthetic response patterns among cultivars. Under salinity, Kumen, Ameristand, and Chenini maintained relatively stable Phi2, with no significant changes over time. In contrast, the remaining cultivars exhibited significant declines, particularly MSI037 (−31.11%), followed by Salado (−22.22%), MSI038 (−19.57%), and MSI036 (−19.51%).

### 3.6. Integrative Trait Network Analysis Links Temporal Chlorophyll Fluorescence Dynamics and Foliar Na□ to Biomass Maintenance

To broadly map multi-trait interaction networks without a priori functional assumptions, pairwise Spearman correlations were performed across the complete 52-variable dataset (1,378 unique pairwise comparisons; Fig. S4). Given the extreme statistical penalty inherent to Benjamini-Hochberg FDR adjustments over high-dimensional exploratory matrices, nominally significant correlations (126 pairs, P < 0.05) were utilized strictly for pattern discovery rather than univariate hypothesis testing. To resolve biologically relevant trait clusters, a targeted correlation sub-matrix (Fig. 4) was constructed focusing on 12 core physiological traits that demonstrated consistent covariance and strong alignment with principal component axes.

**Fig. 4.**
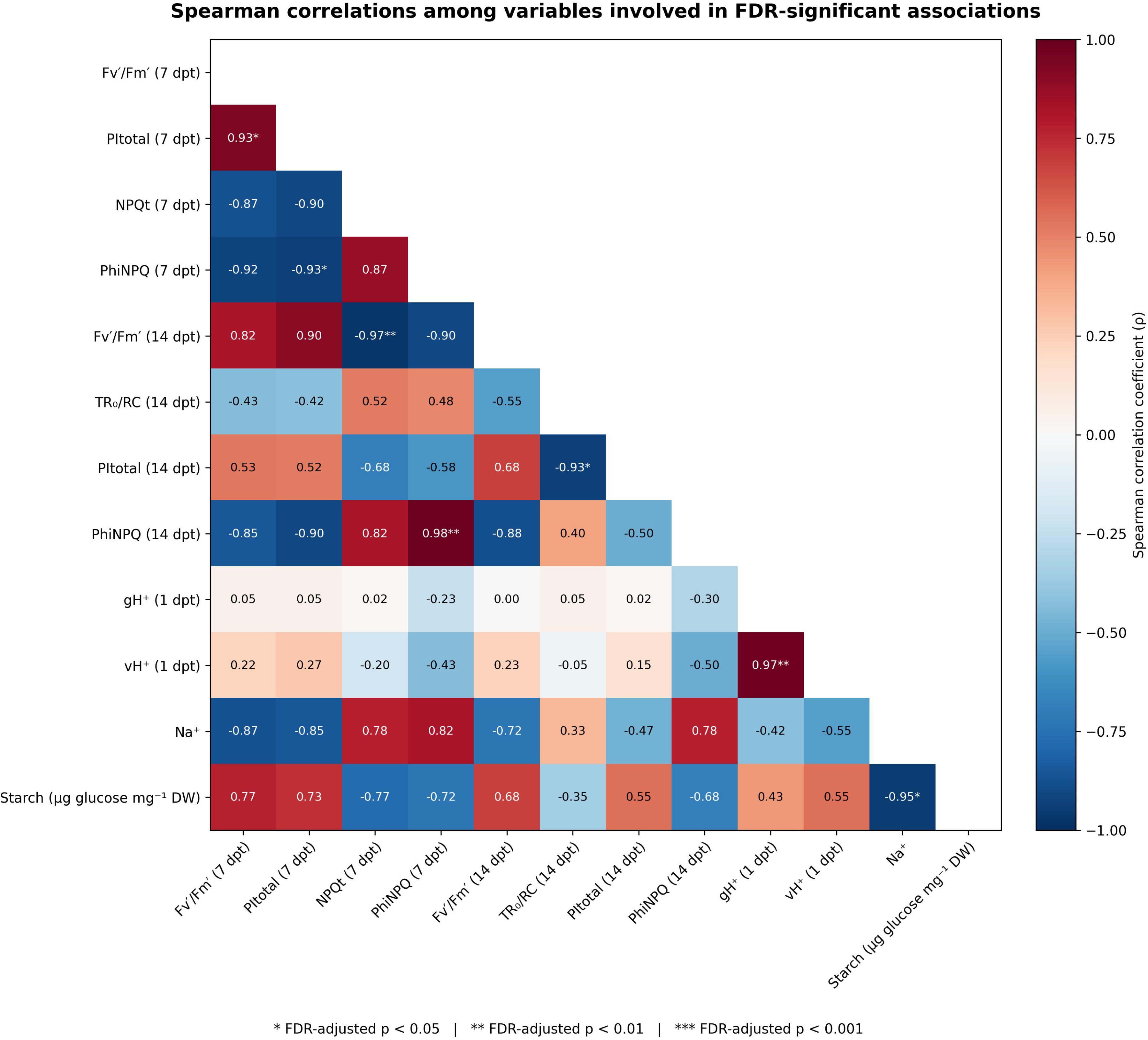
Spearman correlation matrix of variables involved in the most robust associations detected across cultivars. Spearman correlation matrix showing the relationships among the 12 variables involved in the seven pairwise associations that remained significant after Benjamini–Hochberg FDR correction in the complete correlation analysis comprising 52 measured traits and shoot biomass retention (53 variables in total). The matrix includes relationships among the selected variables, allowing visualization of the broader correlation structure. Correlation coefficients (ρ) are represented by a diverging color scale centered at zero, with blue indicating negative and red indicating positive associations. Significance annotations correspond to FDR-adjusted *P* values.

Among the strongest observed associations, several involved photochemical performance and energy dissipation. At 7 dpt, Fv’/Fm’ was positively associated with PI_total_ (ρ = 0.93), whereas PhiNPQ was negatively associated with PI_total_ (ρ = −0.93). NPQt at 7 dpt was strongly negatively associated with Fv’/Fm’ at 14 dpt (ρ = −0.97). The strongest positive association was observed between PhiNPQ at 7 and 14 dpt (ρ = 0.983), suggesting that cultivar-specific patterns of energy dissipation were maintained across these sampling times. At 14 dpt, PI_total_ was negatively associated with TRo/RC (ρ = −0.93). Among early responses, gH□and vH□ at 1 dpt were strongly positively associated (ρ = 0.97). In addition, foliar Na□ concentration was negatively associated with starch content (ρ = −0.95).

Associations between ChlF responses at 1, 7, and 14 dpt and final shoot biomass retention were also examined. Several relationships showed nominal associations with biomass retention; however, these did not remain significant after correction for multiple comparisons across the complete correlation matrix. At 1 dpt, PhiNPQ showed a positive association with biomass retention (ρ = 0.72). At 7 dpt, PI_total_ and Fv’/Fm’ showed strong positive associations with biomass retention, whereas PhiNPQ and PhiNO showed negative associations (Fig. 5). NPQt and gH□also displayed negative trends. At 14 dpt, Fv’/Fm’ and Phi2 showed positive associations with biomass retention, while PhiNPQ remained negatively associated.

**Fig. 5.**
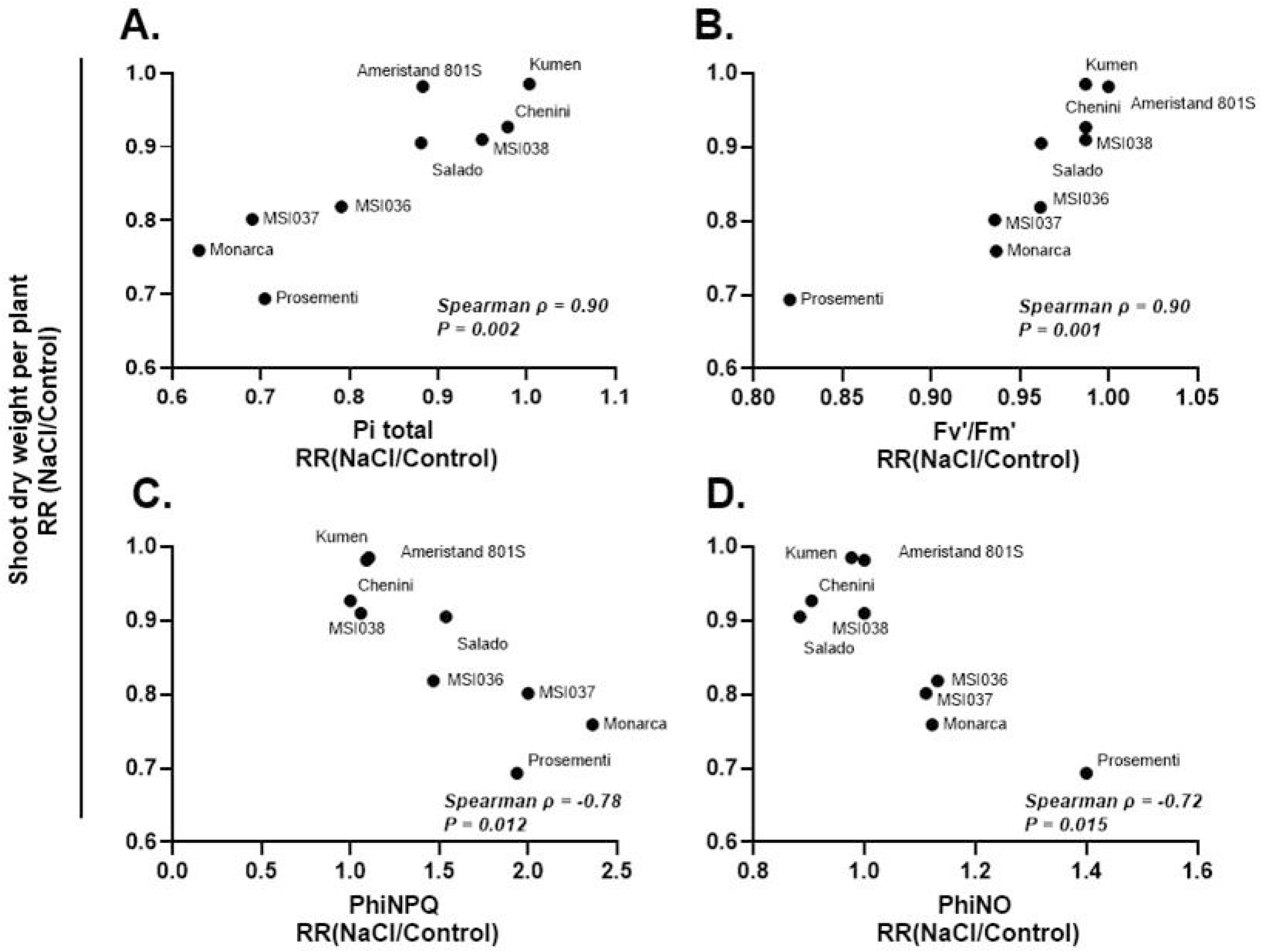
Correlation between photochemical traits at 7 dpt and final shoot biomass under salinity. Spearman rank correlations between the response ratio (RR; NaCl/Control) of shoot dry weight per plant and the RR of (A) PItotal, (B) Fv’/Fm’, (C) PhiNPQ, and (D) PhiNO measured at 7 dpt. Each point represents one alfalfa cultivar (n = 9). Response ratios were calculated as the mean value under 200 mM NaCl divided by the corresponding control mean for each cultivar. Spearman’s ρ and associated *P*-values are indicated in each panel.

Notably, the direction of the PhiNPQ–biomass retention relationship changed over time, from positive at 1 dpt to negative at 7 and 14 dpt. This temporal pattern may reflect differences in the role of energy dissipation during the progression of salt stress. Early increases in PhiNPQ could be associated with an initial photoprotective response, whereas sustained or elevated energy dissipation at later stages may indicate persistent stress and a greater limitation in maintaining photosynthetic performance. Nevertheless, these relationships should be interpreted as exploratory associations rather than evidence of a causal mechanism. Consequently, while individual univariate correlations within the high-dimensional matrix serve as exploratory associations, the definitive validation of these trait relationships requires dimension-reducing multivariate models that bypass pairwise multiple-testing constraints.

### 3.7. Multivariate Trait Integration and Principal Component Analysis of Alfalfa Cultivar Responses to Salinity

Integration of the measured traits revealed distinct cultivar-specific response profiles under salinity (Fig. 6). Cultivars with greater biomass retention, particularly Kumen, Ameristand 801S, Chenini, and MSI038, generally exhibited smaller or transient alterations in photochemical performance and electron transport, together with comparatively limited changes in growth traits. In contrast, cultivars with lower biomass retention, particularly Monarca and Prosementi, showed broader and more persistent reductions in photochemical performance, accompanied by stronger changes in non-photochemical energy dissipation and growth-related traits. Salado, MSI036, and MSI037 displayed intermediate or contrasting response patterns depending on the physiological process considered. Variation in ion accumulation, non-structural carbohydrate pools, and forage-quality traits further demonstrated that cultivar responses to salinity involved changes across multiple physiological levels. Overall, the integrated profiles were consistent with the cultivar ranking based on final biomass retention.

**Fig. 6.**
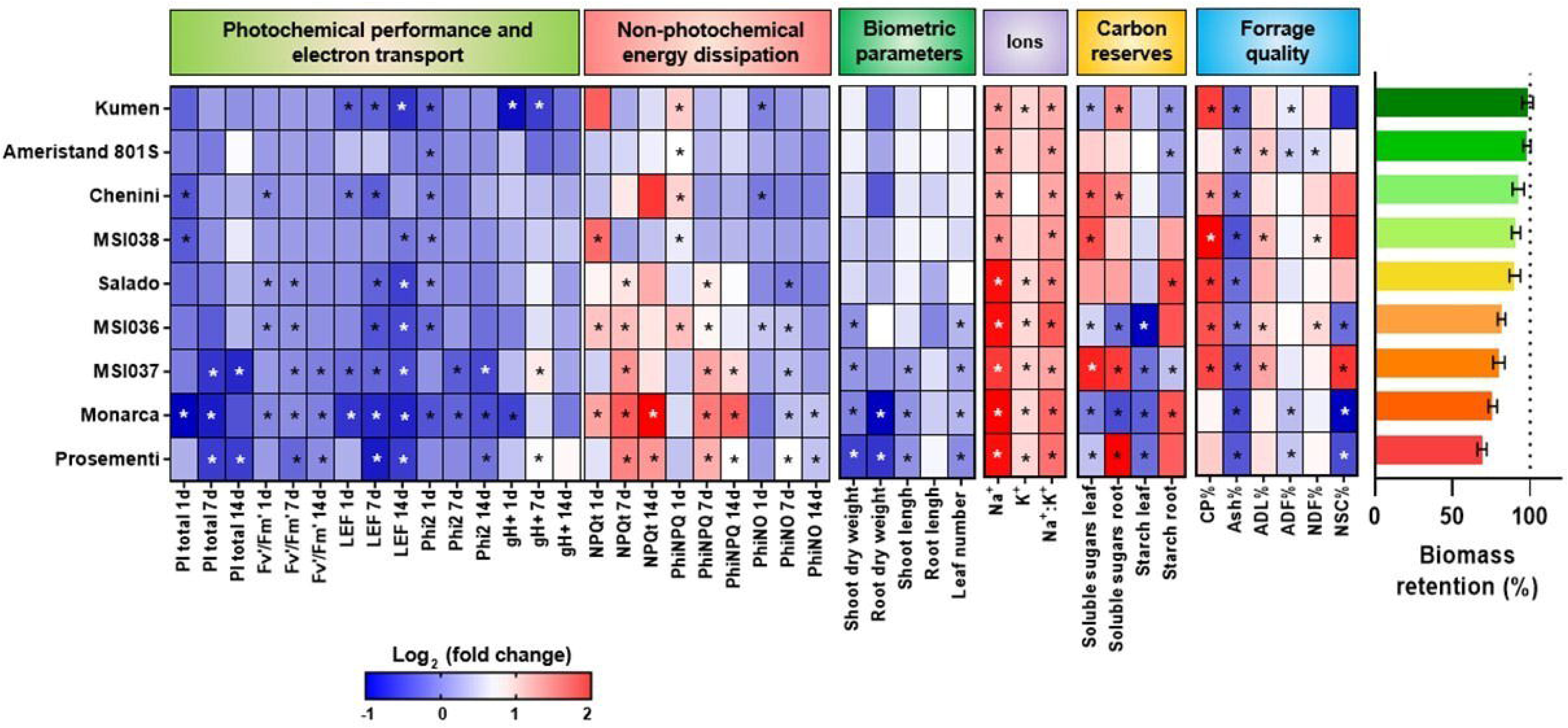
Integrated physiological responses of nine alfalfa cultivars to salinity. Heatmap summarizing cultivar responses to 200 mM NaCl across photochemical performance and electron transport, non-photochemical energy dissipation, growth traits, ion concentrations, non-structural carbohydrates, and forage-quality parameters. Heatmap values represent log fold changes (NaCl/Control), with blue indicating decreases and red indicating increases relative to the corresponding control. Photochemical parameters are shown at 1, 7, and 14 dpt where applicable. Horizontal bars indicate final shoot biomass retention under salinity, expressed as a percentage of the corresponding control. Cultivars are ordered from highest to lowest biomass retention to facilitate comparison between integrated physiological responses and salinity tolerance. Asterisks indicate significant differences between NaCl-treated plants and their corresponding controls within each cultivar and sampling time, as determined by ANOVA followed by Tukey’s post hoc test (P < 0.05).

To further resolve the multivariate structure of these responses, a principal component analysis (PCA) was performed using the salinity responses of the 52 measured physiological, photochemical, ionic, and biochemical variables (Fig. 7A). The first principal component (PC1) explained 34.1% of the total variance, while PC2 explained 21.1%, together accounting for 55.2% of the total variation. Cultivars were distributed broadly across the PC1–PC2 space, indicating substantial differences in their integrated responses to salinity. Kumen and Ameristand 801S, which exhibited the highest biomass retention, were positioned toward the positive PC1 region, whereas Monarca and Prosementi, which showed lower biomass retention, were located toward the negative side of PC1. MSI038 also showed a relatively high positive PC1 score, while the remaining cultivars occupied intermediate positions. This distribution indicates that the principal multivariate axis captured a broad, multi-trait physiological gradient underlying salinity response, which fundamentally aligns with and explains overall biomass preservation.

**Fig. 7.**
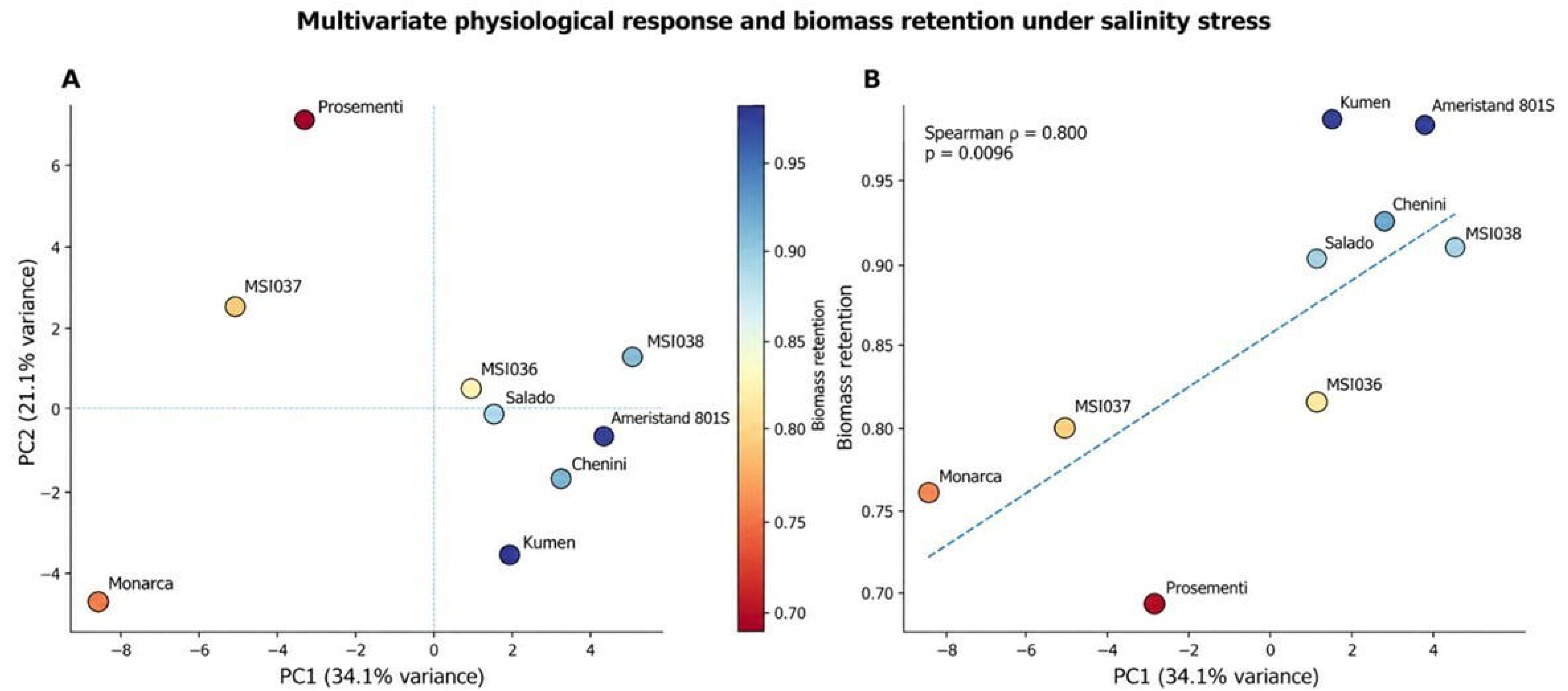
Principal component analysis of cultivar-level salinity responses and association with biomass retention in alfalfa. **(A)** Principal component analysis (PCA) of 52 physiological, photochemical, ionic, and biochemical variables across nine alfalfa cultivars. Salinity responses were expressed as log fold changes (NaCl/Control), and variables were standardized before analysis. PC1 and PC2 explained 34.1% and 21.1% of the total variance, respectively, together accounting for 55.2%. Cultivar positions reflect similarities and differences in their integrated responses to salinity. **(B)** Relationship between PC1 scores and final shoot biomass retention, calculated as shoot dry weight under 200 mM NaCl divided by the corresponding control value. The association was assessed using Spearman’s rank correlation coefficient (ρ). Each point represents one cultivar (*n* = 9). The fitted line is included to visualize the overall trend.

PC1 scores were positively associated with final shoot biomass retention across cultivars (Spearman’s ρ = 0.800, *P* = 0.0096, *n* = 9; Fig. 7B), with higher scores corresponding to greater biomass retention under salinity. This relationship linked the multivariate physiological response profiles to cultivar differences in biomass maintenance.

## 4. Discussion

This study provides new insights into the physiological basis of intraspecific variation in salt tolerance in alfalfa and its implications for forage quality. The cultivar responses characterized through shoot and root traits under hydroponic conditions were largely consistent with field performance previously reported by Pecetti *et al*. (2024) in saline environments in Argentina and Tunisia. This agreement supports the suitability of the hydroponic system as a rapid phenotyping approach for characterizing genotypic variation in salt tolerance and identifying promising candidates for breeding programs. Specifically, our results corroborate the superior salt tolerance of cv. Kumen (previously evaluated as experimental line SISA14; Cornacchione & Suarez, 2017). An exception to this field-hydroponic correlation was cv. MSI037. Although MSI037 exhibited consistently high long-term biomass yield in saline fields, it did not rank among the most tolerant cultivars in our acute stress assay. This discrepancy suggests that short-term physiological responses may not fully capture cultivar resilience under chronic field salinity. Nevertheless, rather than segregating cultivars into strictly tolerant or sensitive categories, our findings reveal a phenotypic continuum in which different cultivars deploy distinct combinations of adaptive traits. For instance, despite its lower overall biomass ranking in this study, MSI037 exhibited K□ accumulation coupled with enhanced soluble sugar accumulation in both leaves and roots likely supporting osmotic adjustment. Furthermore, maintaining K□ homeostasis is proposed as a critical determinant of salinity tolerance in alfalfa, particularly under severe NaCl stress (150–200 mM), where ionic imbalances drive severe growth inhibition and loss of vegetative vigor (Wang *et al*., 2026). Ultimately, these genotype-specific responses underscore the mechanistic diversity of alfalfa salt adaptation, demonstrating that distinct physiological strategies can optimize performance under stress without necessarily converging on a single, broad-spectrum tolerance phenotype.

Our results indicate that salt tolerance in alfalfa is not determined by the absence or magnitude of the initial stress response, but rather by the capacity to sustain physiological functioning during prolonged salt exposure. Although all cultivars exhibited physiological adjustments in response to salinity, more tolerant cultivars, including Ameristand 801S, MSI038, Salado, Kumen, and Chenini, better maintained growth and biomass accumulation, whereas Prosementi and Monarca showed more pronounced growth inhibition. These contrasting phenotypic responses were accompanied by differences in photochemical stability, ion homeostasis, and organ-specific carbohydrate dynamics. Collectively, our findings suggest that intraspecific variation in alfalfa salt tolerance arises from the coordinated contribution of photosynthetic performance, ionic regulation, and carbohydrate metabolism (Daud *et al*., 2026).

Beyond biomass preservation, our results revealed substantial cultivar-specific changes in forage nutritional quality under salinity. Salinity can reduce biomass yield even when forage quality is maintained or modestly improved (Anderson *et al*., 2023; Díaz *et al*., 2018; Pérez-Reverón *et al*., 2024). CP content tended to increase under salt stress, consistent with previous reports under mild salinity (Anderson *et al*., 2023; Ferreira *et al*., 2015). Such increases may partly result from a concentration effect associated with reduced biomass accumulation and changes in leaf-to-stem allocation, given the generally higher protein content of leaves compared with stems (Pecetti *et al*., 2017). However, CP content did not increase significantly in Ameristand 801S, Prosementi, or Monarca, despite their contrasting growth responses. While Ameristand 801S largely maintained growth under salinity, Prosementi and Monarca exhibited fewer leaves and shorter stems, indicating that changes in plant architecture did not necessarily translate into increased CP concentration. Changes in structural fiber fractions also varied among cultivars, with potential implications for forage digestibility and intake (Van Soest, 1994). The significant increases in NDF and lignin (ADL) observed in MSI036 and MSI038 suggest cultivar-specific alterations in cell wall composition that could negatively affect forage digestibility (Le Gall *et al*., 2015; Vanholme *et al*., 2019). Conversely, ADF decreased in Ameristand 801S, Monarca, and Prosementi, while NDF also declined in Ameristand 801S. These contrasting responses indicate that changes in individual nutritional components do not necessarily reflect a consistent improvement or deterioration in overall forage quality. Furthermore, lower NSC content in the less tolerant cultivars (MSI036, MSI037, Monarca, and Prosementi) may represent an additional nutritional and metabolic consequence of salt stress. Collectively, these findings indicate that increased CP content alone should not be interpreted as an overall improvement in forage nutritional quality. Rather, salinity induced cultivar-specific changes in protein, fiber, and non-structural carbohydrate fractions, which may have different implications for forage utilization. Therefore, cultivar selection for saline environments should integrate salt tolerance, biomass production, and a comprehensive assessment of forage nutritional quality to identify genotypes capable of sustaining both productivity and nutritional value under salinity stress.

Comparative evaluations of plant germplasm have shown that salt-tolerant genotypes tend to limit Na□ accumulation and maintain higher foliar K□ concentrations, thereby preserving ionic homeostasis under salinity (Wang et al., 2026; Feng et al., 2025; Yin et al., 2023; Li et al., 2022). This pattern is consistent with findings reported in different crops, including pearl millet (*Pennisetum glaucum* L.), where genotypic variation in ionic regulation has been associated with differences in salinity tolerance (Rashid et al., 2026). Similarly, studies investigating stress mitigation through exogenous silicon application have highlighted the importance of maintaining a favorable leaf K□ /Na□ratio as a component of improved salt tolerance (Meng et al., 2020). Consistent with these established mechanisms, the differential capacity for photochemical stability observed among cultivars studied here was intimately linked to their distinct leaf ion accumulation profiles. Although foliar Na□ increased across all evaluated cultivars, the excessive Na□ load accumulated by specific cultivars likely drove the persistent photochemical impairment recorded at 14 dpt. Because toxic Na□ accumulation is known to directly disrupt chloroplast function and compromise thylakoid membrane integrity (Deinlein *et al*., 2014; van Zelm *et al*., 2020), this ion-induced damage provides a clear mechanistic basis for the sustained decline in PSII efficiency observed in the less tolerant lines. Our results suggest that salt tolerance was more closely associated with the maintenance of relative ionic balance (Na^+^/K^+^ ratio) than with K□ accumulation alone (Chen *et al*. 2007; Shabala & Cuin 2008). In this regard, Monarca and Prosementi exhibited substantial increases in K□ concentration, but their inability to restrict Na□ accumulation resulted in unfavorable Na^+^/K^+^ ratios. In contrast, the exceptionally high K□ accumulation observed in MSI037 appeared sufficient to partially offset Na□ accumulation and maintain an ionic balance. These differences in ionic homeostasis were accompanied by contrasting carbohydrate profiles. Kumen and Chenini maintained or increased root soluble sugars while preserving growth, whereas Monarca and MSI036 showed marked reductions in root soluble sugars and biomass under salinity. Moreover, the reduction in root starch observed in Kumen, Chenini, and Ameristand 801S may reflect mobilization of carbohydrate reserves under stress (Thalmann & Santelia 2017; Dong & Beckles 2019). Together, these results suggest that salt tolerance involves coordinated adjustments in ionic homeostasis and carbohydrate metabolism, potentially contributing to osmotic adjustment and the maintenance of growth under salinity (Krasensky & Jonak 2012).

Previous multivariate analyses in *Arabidopsis thaliana* identified non-photochemical quenching as an important contributor to early salt-stress responses, whereas quantum yield was more strongly associated with long-term performance (Awlia *et al*., 2016). Consistent with an early perturbation of photosynthetic performance, our time-resolved ChlF measurements revealed reductions in Phi2, ETo/RC, and LEF at 1 dpt in several alfalfa cultivars. Although most of these early photochemical changes were not significantly associated with final biomass, PhiNPQ measured at 1 dpt showed a positive correlation with subsequent biomass accumulation. This association is consistent with a potential photoprotective role of regulated energy dissipation during the initial phase of salt stress, as proposed for photosynthetic acclimation mechanisms (Ruban 2016). By 7 dpt, clearer associations emerged between photochemical performance and final biomass (Fig. 5). At this stage, higher PI_total_ and Fv’/Fm’ values were positively associated with final biomass, indicating that the preservation of photosynthetic performance was linked to greater biomass retention under salinity. Conversely, less tolerant cultivars showed greater photochemical impairment, accompanied by sustained increases in regulated and non-regulated energy dissipation, as indicated by elevated NPQt, PhiNPQ, and PhiNO. Collectively, these findings demonstrate that dynamic ChlF parameters—specifically intermediate _total_ (7 dpt) and late /(14 dpt)—do not act as isolated univariate drivers, but as core functional nodes within a multivariate response network. Their strong alignment with the biomass-predictive PC1 axis ( 0.800, 0.0096) validates these non-destructive photochemical traits as robust candidate indicators for salinity tolerance screening (Feng *et al*. 2025).

Although this study provides insights into the interplay among photosynthetic performance, ionic homeostasis, and carbon partitioning in alfalfa under salinity, it was conducted under controlled experimental conditions. As expected, the cultivar ranking for salt tolerance observed here did not fully match that reported in multi-year field experiments (Pecetti *et al*., 2024), highlighting the influence of environmental conditions on genotype performance. Therefore, further studies under field conditions, where salinity and other environmental factors fluctuate, are needed to assess the consistency of these responses. In particular, the potential of dynamic ChlF traits, including PI_total_, LEF, and Fv’/Fm’, as indicators of salt tolerance should be evaluated across larger and more diverse germplasm collections under both managed and naturally saline conditions.

Overall, our findings indicate that variation in alfalfa salt tolerance is associated with the coordinated maintenance of photochemical performance, ionic balance, and organ-specific carbohydrate adjustments over the course of stress exposure. Ameristand 801S, MSI038, Chenini, and Kumen displayed the most robust short-term response profiles, combining better biomass maintenance with greater photochemical stability. Our integrative findings demonstrate that intermediate photochemical stability (7 dpt) provides a reliable physiological signature, distinguishing tolerant germplasm prior to the onset of severe visual symptoms. Integrating these physiological indicators into broader phenotyping strategies could improve the identification and selection of alfalfa germplasm with enhanced adaptation to saline environments, while field validation remains an essential step toward their practical application in breeding programs.

## Abbreviations

ADF: acid detergent fiber
ADL: acid detergent lignin
ChlF: chlorophyll *a* fluorescence
CP: crude protein
Dpt: days post treatment
DW: dry weight
NDF: neutral detergent fiber

## Supplementary data

Fig. S1. Experimental set-up.

Fig. S2. Contrasting shoot phenotypes of alfalfa cultivars under salinity.

Fig. S3. Temporal photochemical responses of nine alfalfa cultivars under salinity.

Fig. S4. Correlation matrix of physiological, photochemical, ionic and biochemical variables.

Table S1. Chlorophyll fluorescence parameters.

Table S2. Effect of salt stress on leaf ion accumulation and Na^+^/K^+^ ratio in alfalfa cultivars.

Table S3. Organ-specific sugar and starch partitioning under salinity.

## Acknowledgments

We thank Daniela Ortiz (INTA Anguil) for generating the reference data, performing the wet chemistry analyses, and developing the NIRS calibration models for alfalfa. We also thank Mónica Cornacchione, Ariel Odorizzi, and Luciano Pecetti for providing alfalfa seeds, and Paola Suarez for technical assistance.

## Author contributions

MR conceptualization. GB and MR designed and performed the experiments. GB, MC methodology. GB, MC, FL, PA and MR formal analysis. GB and MR created and edited the final Figures. GB, MR writing -original draft. GB, MC, FL, PA, and MR writing - review & editing. MR: supervision; MR: funding acquisition.

## Conflict of interest

No conflict of interest declared.

## Funding

This work was supported by Agencia Nacional de Promoción Científica y Tecnológica (ANPCyT, grant PICT2021-00229), Consejo Nacional de Investigaciones Científicas y Técnicas (CONICET grant PUE-UDEA2018), Instituto Nacional de Tecnología Agropecuaria (grant INTA-2023-PD-I084; INTA-2023-PD-I100).

## Data availability

The data that support the findings of this study are available from the corresponding author upon reasonable request

## Notes

### Competing Interest Statement

The authors have declared no competing interest.

