## Supplemental File for "Integrating Biometric, Metabolic, and Photochemical Traits to Identify Candidate Physiological Indicators of Salinity Tolerance and Forage Quality in Alfalfa Cultivars"

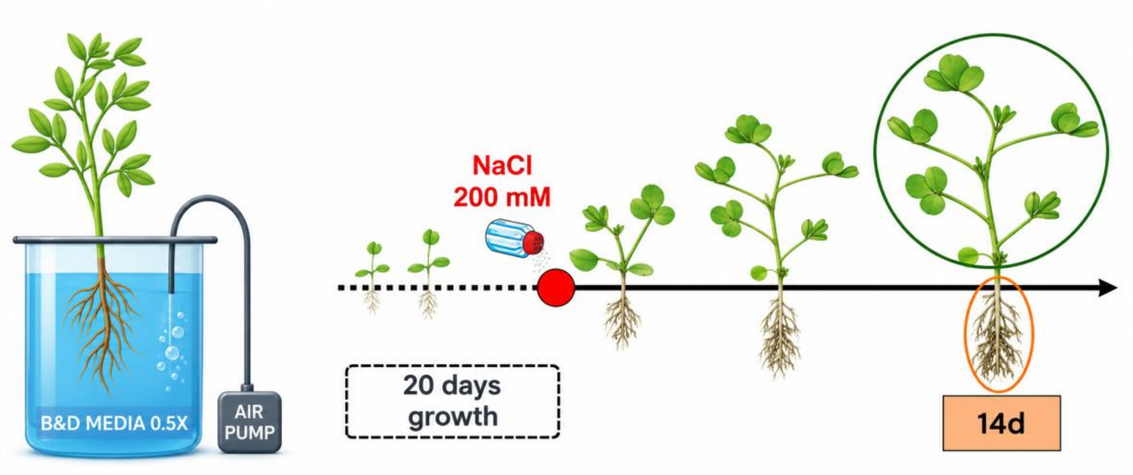


**Figure S1.** Hydroponic growth system and timeline of treatment application and sampling for alfalfa cultivars.

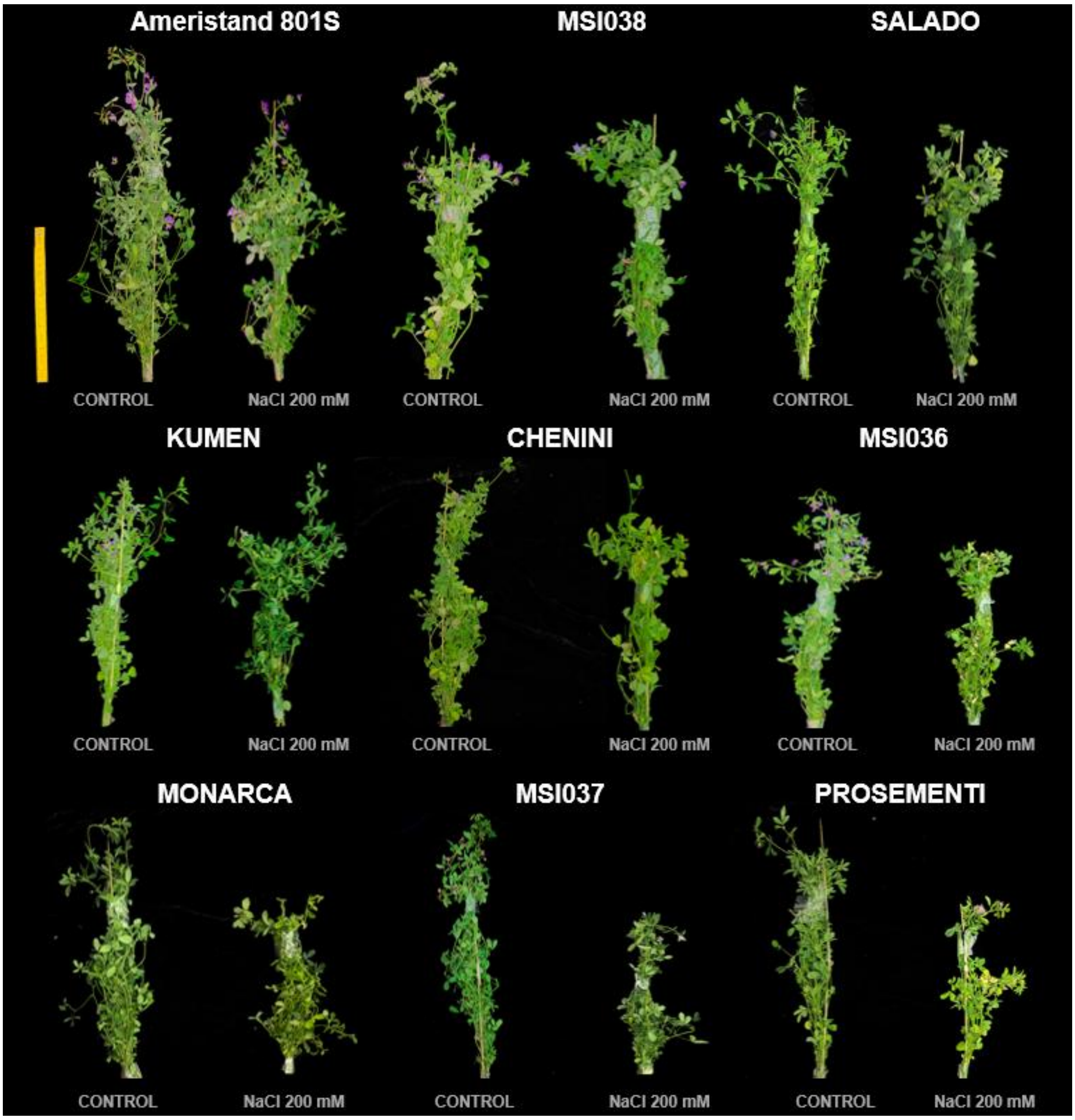


**Figure S2. Contrasting shoot phenotypes of alfalfa cultivars under salinity.** Representative shoots of nine alfalfa cultivars (Ameristand, MSI038, Salado, Kumen, Chenini, MSI036, Monarca, MSI037, and Prosementi) after 14 days under control conditions and 200 mM NaCl treatment. Each bundle comprises 20 plants. For each cultivar, control plants are shown on the left and salt-treated plants on the right.

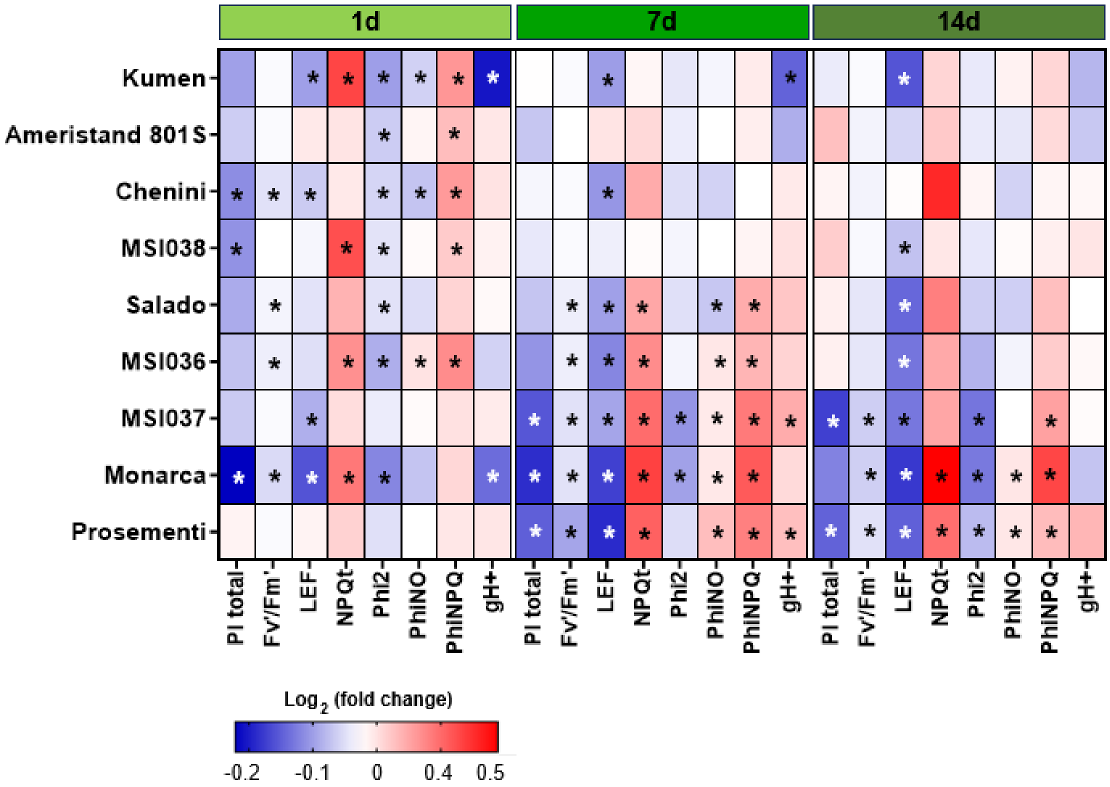


**Figure S3. Temporal photochemical responses of nine alfalfa cultivars under salinity.** Heatmap showing changes in PItotal, Fv/Fm, LEF, NPQ, Phi2, PhiNO, PhiNPQ, and $gH^+$ at 1, 7, and 14 days post treatment (dpt) with 200 mM NaCl. Values represent $log_2$ fold changes calculated relative to the corresponding controls (NaCl/Control). Colors indicate the direction and magnitude of the responses according to the scale shown. Asterisks indicate significant differences between NaCl-treated plants and their respective controls at each sampling time ($P < 0.05$).

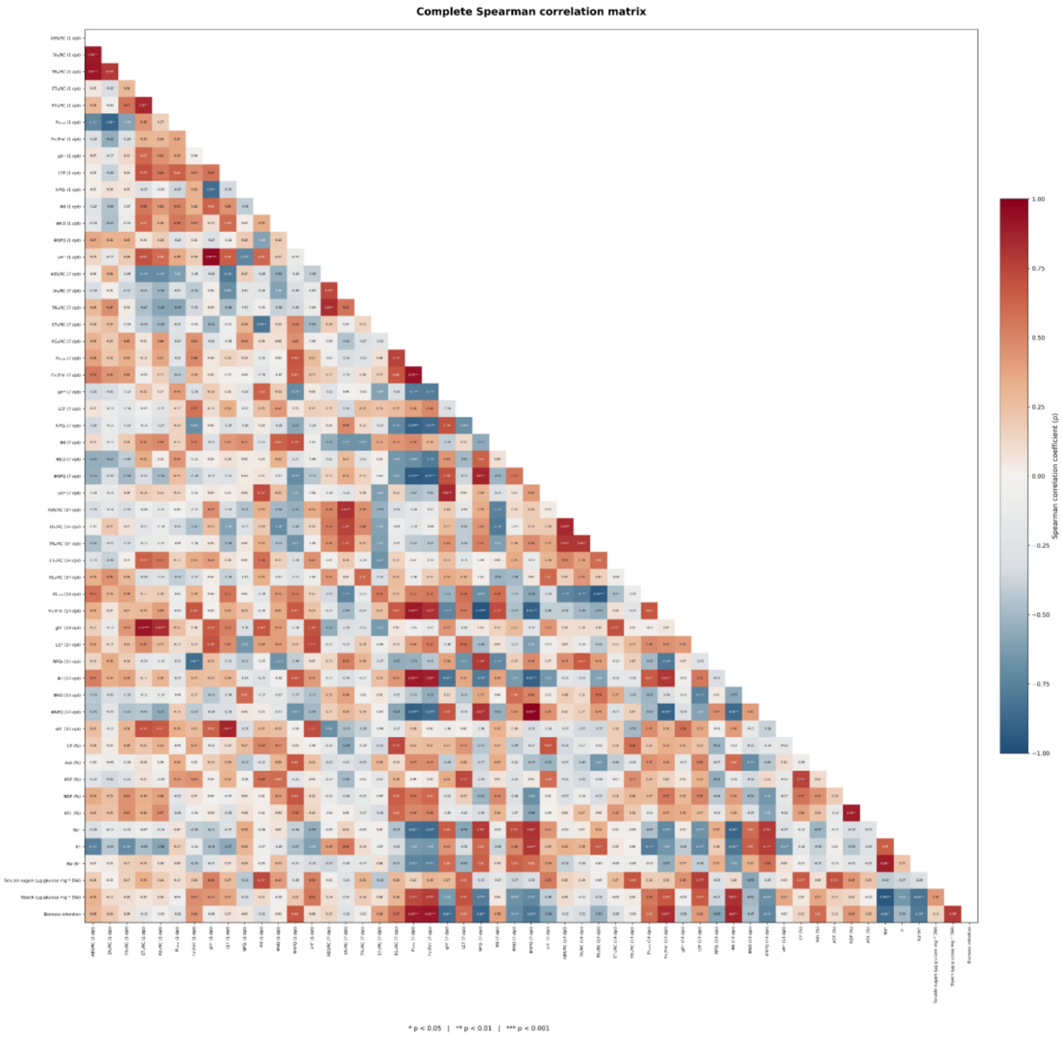


**Figure S4. Complete Spearman correlation matrix of physiological, photochemical, ionic and biochemical variables.** Complete Spearman rank correlation matrix including the 52 measured variables and biomass retention across the nine alfalfa cultivars. Correlations were calculated at the cultivar level using the corresponding salinity responses. The color scale represents Spearman's correlation coefficient (ρ), ranging from −1 to +1 and centered at zero. Blue indicates negative associations, whereas red indicates positive associations. Significance annotations indicate nominal *P* values (*P* < 0.05, P < 0.01, *P* < 0.001). This matrix is provided as exploratory supplementary material to visualize the complete correlation structure among the measured traits.

**Supplementary Table S1.**

| Chlf Parameter. | Description. |
|---|---|
| **Fv'/Fm' (Maximal photochemical efficiency of PSII)** | Key indicator of the photochemical efficiency of PSII and is often used as a measure of plant health and stress. |
| **Phi2 (Effective quantum yield of PSII photochemistry)** | Fraction of excitation energy absorbed by PSII used for photochemistry under light-adapted conditions. |
| **Abs/rc (Absorption per reaction center)** | Represents the amount of light absorbed by each PSII reaction center, helping to evaluate the light-harvesting efficiency of PSII. |
| **Dio/rc (Dissipated energy flux per active PSII reaction center)** | Excitation energy dissipated rather than trapped for photochemistry, expressed per active PSII reaction centre. |
| **Tro/rc (Trapped excitation energy flux per active PSII reaction center)** | Efficiency of the photosynthetic apparatus in converting absorbed light into energy stored in chemical form. |
| **Eto/rc (Electron transport per reaction center)** | Rate of electron transport per PSII reaction center, providing insights into the photosynthetic capacity of the plant. |
| **Reo/rc (Electron flux reducing terminal PSI acceptors per active PSII reaction center)** | A measure of how efficiently electron transport occurs per PSII reaction center. |
| **PI total (Total performance index)** | Integrated index of photosynthetic performance from light absorption by PSII to the reduction of terminal electron acceptors on the PSI acceptor side. |
| **LEF (Linear Electron Flow)** | Represents the rate of electron transport through photosystems II and I during photosynthesis, supporting ATP and NADPH production. |
| **Gh+ (Proton Conductivity):** | Conductivity of protons through the thylakoid ATP synthase complex, providing an estimate of ATP synthase activity. |
| **Vh+ (Steady-State Proton Flux):** | Rate of proton translocation across the thylakoid membrane, primarily driven by the photosynthetic electron transport chain. |
| **NPQ (Non-photochemical quenching)** | Dimensionless coefficient describing non-photochemical quenching of chlorophyll fluorescence associated with thermal energy dissipation. |

| | |
|---|---|
| **PhiNO (Non-photochemical fluorescence quenching)** | Fraction of excitation energy absorbed by PSII lost through non-regulated dissipation pathways, including fluorescence and constitutive thermal dissipation. |
| **PhiNPq (Non-photochemical quenching efficiency)** | Fraction of excitation energy absorbed by PSII dissipated through regulated photoprotective processes. |

| Cultivar | Treatment | Soluble sugar leaf (µg glucose $mg^{-1}$ DW) | Starch leaf (µg glucose $mg^{-1}$ DW) | Soluble sugar root (µg glucose $mg^{-1}$ DW) | Starch root (µg glucose $mg^{-1}$ DW) |
|---|---|---|---|---|---|
| Kumen | Control | 0.87 ± 0.07 | 3.34 ± 0.25 | 0.32 ± 0.02 | 2.91 ± 0.1 |
| | NaCl 200 mM | 0.59 ± 0.04 * | 2.44 ± 0.26 | 0.40 ± 0.03 * | 1.8 ± 0.16 * |
| Ameristand 801S | Control | 0.67 ± 0.06 | 2.78 ± 0.33 | 0.5 ± 0.03 | 2.72 ± 0.25 |
| | NaCl 200 mM | 0.69 ± 0.09 | 2.48 ± 0.35 | 0.49 ± 0.04 | 1.77 ± 0.21* |
| Chenini | Control | 0.62 ± 0.05 | 3.14 ± 0.28 | 0.35 ± 0.02 | 2.16 ± 0.17 |
| | NaCl 200 mM | 0.86 ± 0.02 * | 2.66 ± 0.26 | 0.43 ± 0.05 * | 1.37 ± 0.05 * |
| MSI038 | Control | 0.58 ± 0.08 | 3.68 ± 0.69 | 0.44 ± 0.01 | 2.12 ± 0.28 |
| | NaCl 200 mM | 0.85 ± 0.03 * | 2.75 ± 0.5 | 0.46 ± 0.03 | 2.47 ± 0.35 |
| Salado | Control | 0.51 ± 0.02 | 3.01 ± 0.31 | 0.37 ± 0.03 | 1.59 ± 0.3 |
| | NaCl 200 mM | 0.58 ± 0.05 | 2.19 ± 0.15 | 0.42 ± 0.01 | 2.43 ± 0.42 * |
| MSI036 | Control | 0.66 ± 0.03 | 3.32 ± 0.47 | 0.58 ± 0.02 | 0.97 ± 0.05 |

| | | | | | |
|---|---|---|---|---|---|
| | NaCl 200 mM | 0.5 ± 0.06 * | 1.2 ± 0.24 * | 0.31 ± 0.03 * | 1.43 ± 0.19 |
| MSI037 | Control | 0.59 ± 0.07 | 2.83 ± 0.11 | 0.28 ± 0.03 | 1.51 ± 0.12 |
| | NaCl 200 mM | 1.06 ± 0.18 * | 1.53 ± 0.17 * | 0.46 ± 0.05 * | 1.07 ± 0.08 * |
| Monarca | Control | 0.78 ± 0.05 | 4.33 ± 0.34 | 0.54 ± 0.01 | 1.75 ± 0.22 |
| | NaCl 200 mM | 0.44 ± 0.03 * | 2.19 ± 0.31 * | 0.27 ± 0.02 * | 2.57 ± 0.08 * |
| Prosementi | Control | 0.8 ± 0.05 | 3.92 ± 0.3 | 0.27 ± 0.02 | 1.06 ± 0.11 |
| | NaCl 200 mM | 0.57 ± 0.06 * | 2.09 ± 0.35 * | 0.48 ± 0.02 * | 1.52 ± 0.26 |

**Supplementary Table S2. Organ-specific sugar and starch partitioning under salinity.**

Soluble sugars and starch were quantified in leaves and roots of nine alfalfa cultivars grown under control conditions or exposed to 200 mM NaCl. Values represent mean ± SD. Asterisks indicate significant differences between plants under saline treatment and their respective controls, based on three independent experiments with four plants per treatment, grouped into one pool (n=3), analyzed using ANOVA followed by Tukey's test ($p < 0.05$).

| Cultivar | Treatment | $Na^+$ (µg mg⁻¹ DW) | $K^+$ (µg mg⁻¹ DW) | $Na^+/K^+$ |
|---|---|---|---|---|
| Ameristand 801S | Control | 0.03 ± 0.01 | 0.14 ± 0.01 | 0.21 ± 0.05 |
| | NaCl 200 mM | 0.68 ± 0.05 * a | 0.17 ± 0.0033 abcd | 4.17 ± 0.40 * ab |
| MSI038 | Control | 0.03 ± 0.0033 | 0.14 ± 0.01 | 0.20 ± 0.02 |
| | NaCl 200 mM | 0.82 ± 0.04 * a | 0.16 ± 0.01 abc | 5.26 ± 0.58 * abc |
| Salado | Control | 0.01 ± 0.0033 | 0.12 ± 0.01 | 0.13 ± 0.04 |
| | NaCl 200 mM | 0.70 ± 0.05 * a | 0.18 ± 0.0033 * bcd | 3.94 ± 0.34 * ab |

| | | | | |
|---|---|---|---|---|
| Kumen | Control | 0.03 ± 0.01 | 0.11 ± 0.01 | 0.26 ± 0.05 |
| | NaCl 200 mM | 0.69 ± 0.08 * a | 0.19 ± 0.01 * cd | 3.51 ± 0.17 * a |
| Chenini | Control | 0.05 ± 0.01 | 0.13 ± 0.02 | 0.39 ± 0.03 |
| | NaCl 200 mM | 0.87 ± 0.12 * a | 0.12 ± 0.02 a | 7.34 ± 1.25 * bcd |
| MSI036 | Control | 0.02 ± 0.01 | 0.10 ± 0.01 | 0.23 ± 0.07 |
| | NaCl 200 mM | 1.42 ± 0.08 * bc | 0.15 ± 0.01 * abc | 9.30 ± 0.95 * de |
| Monarca | Control | 0.02 ± 0.00 | 0.08 ± 0.01 | 0.31 ± 0.03 |
| | NaCl 200 mM | 1.54 ± 0.04 * c | 0.13 ± 0.00 * ab | 11.57 ± 0.19 * e |
| MSI037 | Control | 0.02 ± 0.00 | 0.09 ± 0.01 | 0.23 ± 0.03 |
| | NaCl 200 mM | 1.04 ± 0.14 * ab | 0.21 ± 0.01 * d | 4.81 ± 0.46 * ab |
| Prosementi | Control | 0.02 ± 0.0033 | 0.10 ± 0.01 | 0.24 ± 0.06 |
| | NaCl 200 mM | 1.40 ± 0.15 * bc | 0.17 ± 0.02 * abcd | 8.37 ± 0.99 * cde |

**Supplementary Table S3. Effect of salt stress on leaf ion accumulation and $Na^+/K^+$ ratio in alfalfa cultivars.**

Leaf $Na^+$ concentration, $K^+$ concentration and $Na^+/K^+$ ratio was determined in nine alfalfa cultivars grown under control conditions and exposed to 200 mM NaCl. Values represent mean ± SD. Asterisks indicate significant differences between plants under saline treatment and their respective controls, based on three independent experiments with three plants per treatment, grouped into one pool (n=3), analyzed using ANOVA followed by Tukey's test ($p < 0.05$). Different lowercase letters indicate significant differences among cultivars under NaCl treatment ($P < 0.05$). Cultivars sharing at least one letter are not significantly different.
